# C-terminal tagging impairs AGO2 function

**DOI:** 10.1101/2023.11.20.567203

**Authors:** Kunal M. Shah, Alex F. F. Crozier, Anika Assaraf, Muzjda Arya, Paul Grevitt, Faraz Mardakheh, Michael J. Plevin, Tyson V. Sharp

## Abstract

A full understanding of RNA silencing requires appropriate molecular biology tools to explore the roles of Argonaute 2 (AGO2) and the RNA-induced silencing complex (RISC). Approaches relying on affinity tagging and antibodies have important limitations that can lead to artificial results. Both the N- and C-terminal domains of AGO2 have been shown to be important for correct activity and yet the consequences of appending tags to either terminus have not been fully investigated. N-terminal tags are frequently used to study AGO2 biology. Recently, an N-terminal *HaloTag-Ago2* fusion was reported and examined in mice. While the versatile HaloTag provided new opportunities to study RISC biology, the tagged construct showed certain activity changes compared to unmodified AGO2. CRISPaint, a new CRISPR-Cas9 technique, permits the creation of endogenous C-terminal tag fusions. We used CRISPaint to generate the first reported recombinant AGO2 construct with a C-terminal tag: an endogenous C-terminal HaloTag fusion to AGO2 (*AGO2^HALO^*) in human (A549) cells. We found that the AGO2^HALO^ fusion protein has a reduced capacity to interac with the key protein binding partner TNRC6A and that the C-terminal HaloTag does not affect cell viability. However, the *AGO2^HALO^* fusion significantly impairs RNA cleavage and RNA silencing activity compared to control cells and reduces nuclear localisation of the fusion protein. Using plasmid constructs and transient transfection, we compared AGO2 tagged with EGFP at the N- or C-terminus in siRNA and miRNA reporter gene assays, and cellular localisation. N-terminally tagged AGO2 functioned and localised similarly to WT untagged AGO2, whereas C-terminally tagged AGO2 was impaired in siRNA and miRNA silencing and exhibited poor nuclear and P-body localisation. We conclude that the fusion of a C-terminal HaloTag to AGO2 is not appropriate for studying AGO2 and RISC. Our results assert the importance of comprehensively validating recombinant tagging strategies to ensure that any experimental results generated do not arise from, or are not obscured by, critical functional defects.

## Introduction

The four human Argonaute proteins (AGO1-4) are key components of the microRNA-induced silencing complex (miRISC), which is critical for the correct regulation of gene expression through miRNA-mediated silencing.^1,2,3,4,5,6,7^ Due to the considerable scientific and clinical interest in RNA silencing, exemplified by the recent Nobel Prizes to Ambrose and Ruvkun, a significant body of work has been performed to develop experimental tools for the study of AGO protein function. However, assay design has been limited by methods that rely on antibodies and affinity/auto fluorescent tags, which can be technically challenging and often lack the sensitivity or specificity required to generate appropriately robust results.

Recombinantly tagging a protein of interest is a widely used strategy in molecular biology and biotechnology to purify, visualise, and manipulate target proteins. To overcome challenges related to antibody specificity and pull-down efficiency attempts to study RISC have often used overexpressed, recombinantly-tagged AGOs. Such approaches, however, carry the risk of generating artefacts. For example, a N-terminal Flag-tagged AGO2 construct resulted in five-fold over-expression compared to endogenous AGO2 levels, altered AGO2 sub-cellular localisation, and resulted in the identification of potentially false binding partners.^8^ Such artefacts might arise from either over-expression of the transgene or the presence of the Flag tag itself. Moreover, it has been shown that recombinantly-tagged, over-expressed proteins can non-specifically localise to processing (P) bodies.^10^ Caution should therefore be applied to any study that uses recombinantly-tagged proteins to show that both AGO and TNRC6 proteins are strongly localised to P bodies.^0^ Consequently, approaches that tag endogenous proteins are often deemed more desirable. However, this too can disrupt normal protein function.^11,12^ In rapidly proliferating cell lines, endogenously tagged AGOs have been shown to elute in complexes of more variable size than native AGOs have, which primarily elute as components of high molecular weight complexes.^13,14^ An endogenous N-terminal EGFP-AGO2 fusion protein was not recognised by a pS387-AGO2-specific antibody in immunofluorescence assays, conceivably due to the tag altering AGO2 conformation and therefore reducing pS387 antibody binding.^15^ Even in cases where endogenous tags have been shown to retain core RISC activity, these tagged proteins should only be used to study known RISC functions under specific experimental conditions and periods of expression. The use of novel tags and tagging strategies in different cell models may affect the target protein to unpredictable extents. Nevertheless, protein tagging can expand our ability to study RNA silencing and RISC proteins, albeit with (manageable) technical biases which are inevitable when probing complex biological systems. Thus, to confidently exploit the potential benefits that recombinant tagging of endogenous proteins offers it is important to ensure that the core functionalities of the proteins of interest are sufficiently retained - primarily through considered design and stringent validation - and that technical biases should be considered and minimised when designing experiments and interpreting results.

The HaloTag protein fusion platform is a modular protein tagging system that can be programmed for different functions, proving a versatile technology.^16,17,18^ The modified 33 kDa haloalkane dehalogenase HaloTag protein can covalently bind to synthetic chloroalkane ligands (collectively known as HaloTag ligands) which comprise a chloroalkane linker attached to a variety of useful molecules, such as fluorescent dyes, affinity handles, or solid surfaces. Covalent bond formation between the protein tag and the chloroalkane linker is high affinity, rapid, specific, and irreversible. This enables efficient purification of HaloTag fusion proteins and means this single genetic construct can have multiple capabilities to comprehensively analyse protein function and interaction. Importantly, an endogenous fusion of HaloTag to the N-terminus of *Ago2* (HaloTag-Ago2) has enabled the high-resolution identification of Ago2-miRNA and -mRNA interactions in mice.^14^

Applying UV crosslinking to cells expressing HaloTag fusion proteins enables efficient identification of RNA targets of RNA-binding proteins.^19^ Of note, HaloTag-Ago2 fusions have been used to investigate the association of Ago proteins with RNA in a process termed Halo-Enhanced Ago2 Pulldown sequencing (HEAP-seq).^14^ The HaloTag ligand conjugated to a resin can be used to pull down HaloTag-Ago2 in cells or tissues expressing the fusion. High-throughput sequencing of RNA extracted from purified AGO-miRNA-mRNA complexes then enables the identification of miRNA-mRNA networks under physiological (and/or pathological) conditions. In addition to bypassing the need for radiolabelling, immunoprecipitation, and gel purification, the covalent nature of the interaction between the HaloTag and its ligand simplifies the isolation of AGO-miRNA-mRNA complexes, removing the intrinsic variability of antibody-based approaches. Moreover, as the strong covalent bond permits more stringent washing, there is less potential for non-specific RNAs to be captured. ^14,20^ Accordingly, HEAP-seq was demonstrated to enable both more sensitive and specific identification of AGO protein-RNA interactions than conventional methods which rely on the use of antibodies to precipitate AGO-containing complexes (i.e. CLIP-seq^21^). Furthermore, the HaloTag labelling capacity can also be used to perform live-cell single-molecule imaging to observe dynamic activity and high-resolution investigation of protein-protein interactions through mass spectrometry.^17,22,23,24^ Endogenous fusion of AGO2 to HaloTag should therefore facilitate a thorough investigation of AGO2/RISC in human cells at physiologically relevant abundancies, in near-native cellular background, and to greater resolution than antibody-based approaches. Future HaloTag:AGO1-4 fusions would also enable comparative analysis of AGO1-4 biology in different contexts, with the mutual HaloTag overcoming the key limitations of CLIP-based methods that result from variable antibody performance.

AGO2 comprises four domains (N-terminal (N), PIWI/Argonaute/Zwille (PAZ), middle (MID), and P-element-induced wimpy testes (PIWI)), each with known roles in RISC formation and RNA silencing.^25,26,27,28,29,30^ When designing a recombinantly-tagged AGO2 protein, it is critical to consider the impact on both N- and C-terminal domains. The PIWI domain is located at the C-terminus of AGO2. This domain contains six loops along the nucleic acid-binding cleft and is similar in function to an RNase H domain, harbouring the catalytic tetrad (DEDH) essential for the slicing activity of AGO2.^30,31,32,33,34,35,36,37^ Mutation of the conserved glutamate residue in the DEDH motif abolishes the ability of AGO2 to induce RNAi.^36^ Many post-translational modifications (PTMs) of AGO2 are found in the C-terminal PIWI domain.^38^ These PTMs impact various aspects of AGO2 function, influencing both RISC and miRNA activity. The C-terminal half of human AGO2 is also responsible for the interaction with TNRC6A, which is crucial for miRNA-dependent silencing function and the localization of AGO2 in cytoplasmic foci.^6^ The N domain of AGO2 is fundamental to RISC functionality, facilitating ATP-independent unwinding of small RNA duplexes, a critical step for RISC assembly in human cells. Both slicer-dependent and slicer-independent unwinding require efficient function of the N domain of AGO2, highlighting its importance in passenger-strand cleavage. Moreover, two specific motifs (residues 44–48 and 134–166) in the N-terminal region of AGO2 are vital for optimal catalytic activity. It is postulated that, due to their proximity to the PIWI domain in the tertiary structure of AGO2, these motifs are essential for the accurate positioning of the guide:target. Alterations within these motifs hinder the activation of RISC and obstruct mRNA cleavage.^38,39,40^ Deletion of the N-terminal, Linker 1 and PAZ domains in *Drosophila* AGO2 resulted in decreased target RNA binding and constitutive activation of the cleavage activity of the PIWI domain.^41^ These studies highlight the collective importance of both the N- and C-termini in the normal function of AGO2 and emphasise the need for consideration and due diligence when adding and validating tags at either terminus, as each may disrupt normal protein function.

We set out to develop human cell lines endogenously expressing AGO2 fused to a reporter tag to enable comprehensive study of AGO2 function and RISC biology in cells relevant to our research interests. Due to the stated versatility of HaloTag technology and advantages over traditional methods used to study AGO complexes and RISC, we sought to generate a AGO2-HaloTag fusion protein. A previous study that used N-terminal fusion of HaloTag with Ago2 reported diminished ability to rescue RNAi in *Ago2*^-/-^ MEFs compared to WT Ago2, de-repression of a subset of highly expressed miRNA targets, and reduced viability of *HaloTag-Ago2* homozygous mice,^14^ indicating some loss of functionality due to the N-terminal HaloTag-Ago2 fusion. A series of functional validation experiments did, however, demonstrate core functionalities were retained after N-terminal HaloTag-Ago2 fusion. Pull-down assays established physical interaction between HaloTag-Ago2 and the core miRISC component Tnrc6a, while size-exclusion chromatography showed co-elution of HaloTag-Ago2 with WT Ago2 in high molecular weight complexes (though, notably, complexes were of more variable size in tagged cells^9^). Encouragingly, Dual-Luciferase reporter experiments using three constructs with well-characterized miRNA binding sites, as well as a sensitive two-colour fluorescent reporter system,^42^ showed no detectable differences in miRNA-mediated repression between *WT* and *Ago2^Halo/Halo^* MEFs. These findings are an example of orthodox tag validation and, overall, indicate endogenous N-terminal fusion of HaloTag to AGO(2) is a valuable model to study miRISC *in vivo* and *in vitro*, despite some reported functional limitations. However, the tagging approach used by Li *et al*. raises important points of consideration when designing tags. N terminal fusion of HaloTag to Ago2 was achieved by insertion of a conditional knock-in allele containing the HaloTag upstream of a loxP-STOP-IRES-FLAG-loxP cassette in mouse ES cells to express Cre recombinase which fuses the HaloTag to the first exon of *Ago2*.^14^ This is a relatively complex method of gene tagging which can be time-consuming and challenging to precisely reproduce. The approach also disrupts the 5ʹ UTR of the *Ago2* gene, risking unknown effects on the regulation of expression of AGO2, which is critical when considering true *in vivo* endogenous functionality. These challenges therefore prompted us to look for alternative methods to generate an endogenous HaloTag fusion to AGO2 in human cells.

CRISPaint is a gene editing system which allows the precise creation of C-terminal tag fusions of endogenously encoded proteins in human cells with high efficiency.^43^ Unlike homology directed repair (HDR)-directed tagging, CRISPaint does not require generation of a site-specific donor template with homology arms but integrates heterologous genetic material via canonical non-homologous end joining (cNHEJ) in CRISPR-Cas9-accessible cellular systems. This reduces the overall time required to obtain genetically tagged cells compared with HDR-directed tagging.

The lack of data on the validity of C-terminal AGO2 tagging, the non-trivial nature of endogenous N-terminal AGO2 tagging using the method used by Li *et al*, the importance of the 5ʹ UTR, and the critical motifs at both the C- and N-termini of AGO2 together make a preferred strategy for generating tagged variants of endogenous AGO2 tags challenging to predict. We therefore employed CRISPaint to create cell lines expressing a C-terminal AGO2-HaloTag fusion protein (AGO2^HALO^) for the future comprehensive study of AGO2 in human cells. Due to the addition of the tag potentially impairing normal protein function and therefore invalidating any future results, as well as the lack of published evidence on the effect of C-terminal tagging of AGO2, we sought to validate the functional activity of endogenous AGO2^HALO^ robustly and diligently against native AGO2 in UnTagged CRISPaint controls lines and *WT* A549 cells.

By interrogating the core functional activities of AGO2 (protein localisation and binding, cleavage, and silencing capacity) in *WT*, UnTagged, and *AGO2^HALO^*cells, we aimed to define the impact of the C-terminal fusion of AGO2 with HaloTag. Our results led us to conclude that *AGO2^HALO^* cells have significantly impaired silencing function compared to control UnTagged and *WT* AGO2 cells, as well as distinct sub-cellular localisation. We therefore do not consider C-terminal AGO2^HALO^ as a suitable model for further study of AGO2 and RISC function. Although we only considered a single configuration for a single type of C-terminal fusion tag of the relatively large (33 kDa) HaloTag to AGO2, comparison of our data with previously published N-terminal HaloTag-AGO2 does indicate that N-terminal tagging is favourable. Following this, we directly compared the effect of N-vs C-terminal tagging on AGO2 function. Addition of an EGFP tag to the C-terminus of AGO2 resulted in a loss of silencing function comparable to complete AGO2 ablation, whereas addition of the same tag at the N-terminus had no such effect with silencing function maintained. Together, these data indicate C-terminal tagging of AGO2 severely impairs normal AGO2 function and is less suitable than N-terminal AGO2 tagging approaches for study of AGO2/RISC in human cells.

## Materials and Methods

### Cell Culture

A549 cells were maintained in DMEM (Sigma, D8437) supplemented with 10% qualified fetal bovine serum (FBS) (Gibco, 10270106), 100 U/mL penicillin, and 100 µg/mL streptomycin (Gibco, 15140122) in a humidified 37 °C incubator and 5% CO_2_. Cells were passaged every 3-4 days during the log phase of growth with Trypsin/EDTA solution (Gibco, R001100) following a PBS wash. To seed an exact number of cells, cell suspension concentration was counted using a Countess II™ Automated Cell Counter (Invitrogen). Parental stocks were obtained from ATCC. All cell cultures were regularly tested for mycoplasma. Post editing, isogenic lines (A549 WT, UT C1, AGO2^HALO^ C5 and AGO2^HALO^ C10) were authenticated by STR profiling at 16 STR loci (***Supp Fig. 1***) using PCR-single-locus-technology (Eurofins Genomics).

**Figure 1.**
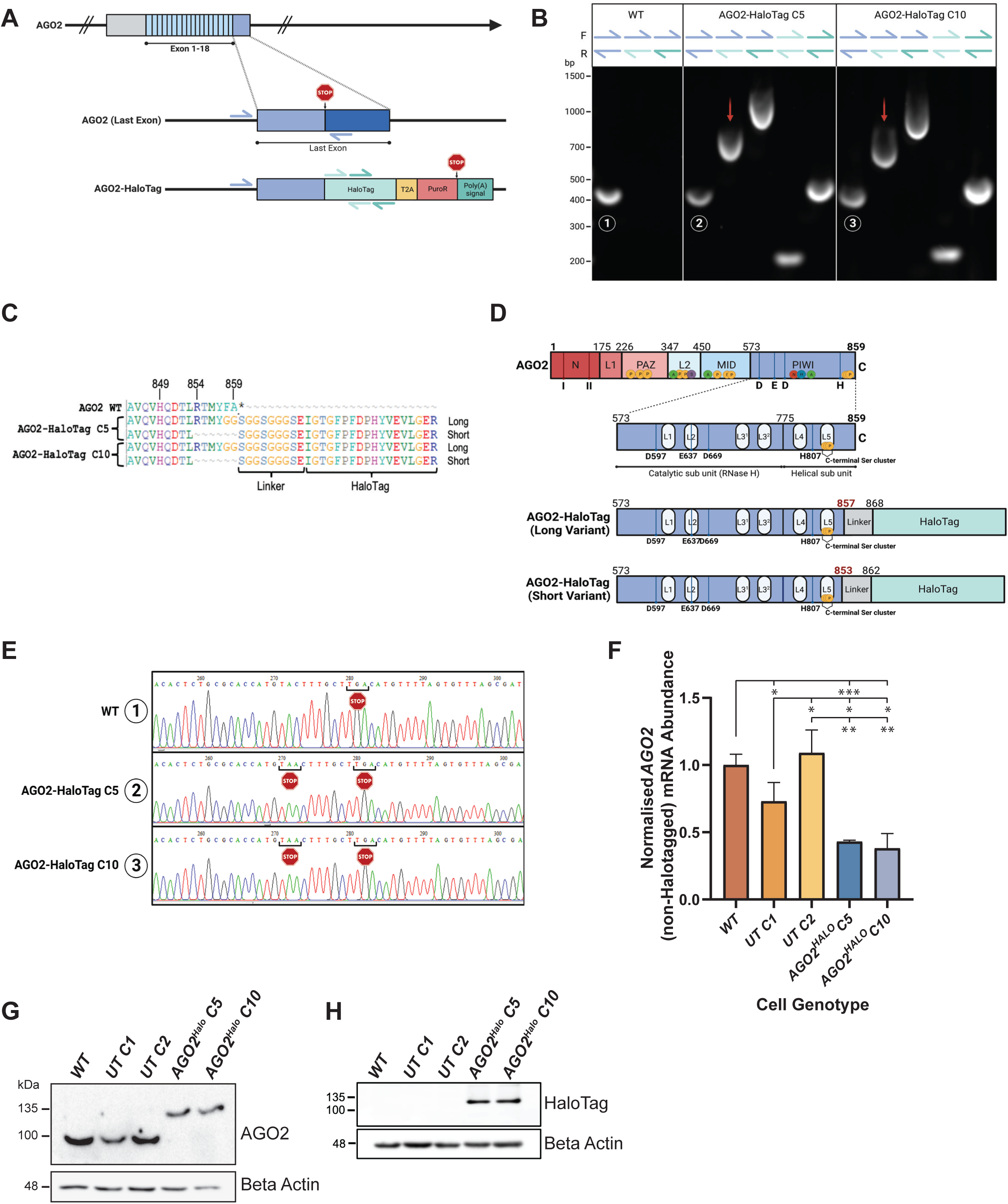
Creation and Genomic Characterisation of AGO2-HaloTag Cell Lines. (A) Schematic of WT AGO2 and the C terminal AGO2-HaloTag fusion (including T2A site, Puromycin resistant (PuroR) and Poly(A) sections) genotype to be generated by CRISPaint editing of A549 cells. Arrows indicate locations of forward and reverse primers designed to confirm editing. Blue = *AGO2* WT last intron Forward and Reverse; Light Teal = HaloTag1 Forward and Reverse; Dark Teal = HaloTag2 Forward and Reverse. (B) Agarose gel loaded with PCR products of A549 WT and two *AGO2-HaloTag* (*AGO2-HaloTag C5* and *AGO2-HaloTag C10*) lines amplified with indicated combinations of *AGO2* WT and *HaloTag* primers, as indicated in (A). Red arrows indicate gDNA containing *HaloTag* sequence which was purified and submitted for sequencing. Circled numbers 1-3 indicate gDNA containing C terminal non-HaloTagged *AGO2* product which was purified and submitted for sequencing. (C) Sequence (generated from TOPO-seq) alignments of *WT* and two *AGO2-HaloTag* clones at the AGO2-HaloTag junction. From several submitted TOPO clones, two variants of *AGO2-HaloTag* (one long and one short) were identified in *AGO2*-*HaloTag* cells. Asterisk (*) indicates STOP codon. (D) Schematic to show known functionally important domains of AGO2, with a focus on C-terminal PIWI domain. CRISPaint mediated AGO2-HaloTag fusion generated a long and a short variant, neither of which resulted in loss of residues known to have functional importance. (E) Chromatograph of C terminal *AGO2* sequence identified in WT, *AGO2-HaloTag C5* and *AGO2-HaloTag C10* cells (Circled numbers 1-3 in (B)) showing the additional and premature STOP codon in both *AGO2-HaloTag* lines. (F) Abundance of non-HaloTagged *AGO2* mRNA transcript in A549 WT, two UnTagged (*UT C1* and *UT C2*) and two *AGO2*-*HaloTag* (*AGO2*-*HaloTag C5* and *AGO2*-*HaloTag C10*) cells. *AGO2* (non-HaloTagged) mRNA abundance normalised to *B Actin* mRNA abundance and made relative to levels in WT cells. Data represent mean ± SEM of experiments; n = 3 (*p ≤ 0.05; **p ≤ 0.01; ***p ≤ 0.001). (G & H) Western blot of whole-cell lysates from A549 WT, two UnTagged (*UT C1* and *UT C2*) and two *AGO2*-*HaloTag* (*AGO2*-*HaloTag C5* and *AGO2*-*HaloTag C10*) cell lines probed with antibodies against AGO2, HaloTag, and Beta Actin.

### CRISPaint Transfection and Puromycin Selection

The CRISPaint gene tagging kit was a gift from Veit Hornung (Addgene kit # 1000000086). To generate a targeting construct for AGO2, we used site-directed mutagenesis to change the gRNA sequence for TUBB to AGO2 gRNA sequence (AACATGTCAAGCAAAGTACA) in the pCas9 mCherry TUBB plasmid. For CRISPaint^43^ transfection, A549 cells were plated in a 96 well plate and, per well, 50 ng pCas9 mCherry AGO2, 50 ng pCas9 mCherry Frame +1 and 100ng pCRISPaint HaloTag-Puro donor plasmids were transfected using ViaFect (Promega E498A). Three days post-transfection cells were sub-cultured into media supplemented with puromycin (0.8 µg/mL; Invivogen, ANT-PR-1). After six days of selection, cells were allowed to recover in media without puromycin for 12 days, followed by another round of puromycin selection for six days. After recovery, the cells were grown out in sufficient quantities to perform Western blotting of whole cell lysates of mixed populations. This was followed by single cell cloning in 96 well plates and once grown in sufficient quantities, individual clones were immunoblotted using the anti-HaloTag antibody (Promega) to screen for the AGO2-HaloTag fusion of approximately 130 kDa molecular weight. Two clones with no tagging of AGO2 (**UnTagged *C1*** and **UnTagged *C2***) and two expressing AGO2-HaloTag (**AGO2^HALO^ C5** and **AGO2^HALO^ C10**) were selected for further evaluation in this study, alongside parental ***A549 WT*** cells. A schematic of the experimental design employed to generate and select *AGO2-HaloTag* (and non-transfected (UnTagged)) CRISPaint clones is provided in ***Fig. 1C***.

### Genomic DNA Sequencing at AGO2-HaloTag Junction

Genomic DNA was prepared by removing the media and lysing the cells in 30 µl of gDNA lysis buffer (0.2 mg per mL proteinase K, 1 mM CaCl_2_, 3 mM MgCl_2_, 1 mM EDTA, 1% Triton X-100 and 10 mM Tris pH 7.5) and then incubating samples at 65 °C for 10 min and at 95 °C for 15 min. The *AGO2-HaloTag* junction region of this genomic DNA was amplified in a 25 µl Phusion Hot Start Flex (M05365S, NEB) PCR reaction as per the manufacturer’s instructions, using specific forward (CAGCTGCTTTTCTGGAAGGG) and reverse (TCTTATGTACCTGACCGACG) primers. Following confirmation of correct PCR amplification on an 0.8% agarose Tris borate EDTA gel, PCR products were TOPO cloned using Zero Blunt™ TOPO™ PCR Cloning Kit for Sequencing (ThermoFisher K287520), transformed into 5-alpha Competent *E. coli* (C2987I, NEB) as per the manufacturer’s protocol for plasmid preparation. Transformed colonies (≥ 5) were grown out in LB medium with 50 µg/mL Kanamycin and then pelleted before plasmids were isolated using the Monarch Plasmid Miniprep Kit (NEB, T1010L). Plasmids were sequenced (Sanger) from the T7 priming site of Blunt II-TOPO vector. Sequencing alignment was performed on BioEdit software.

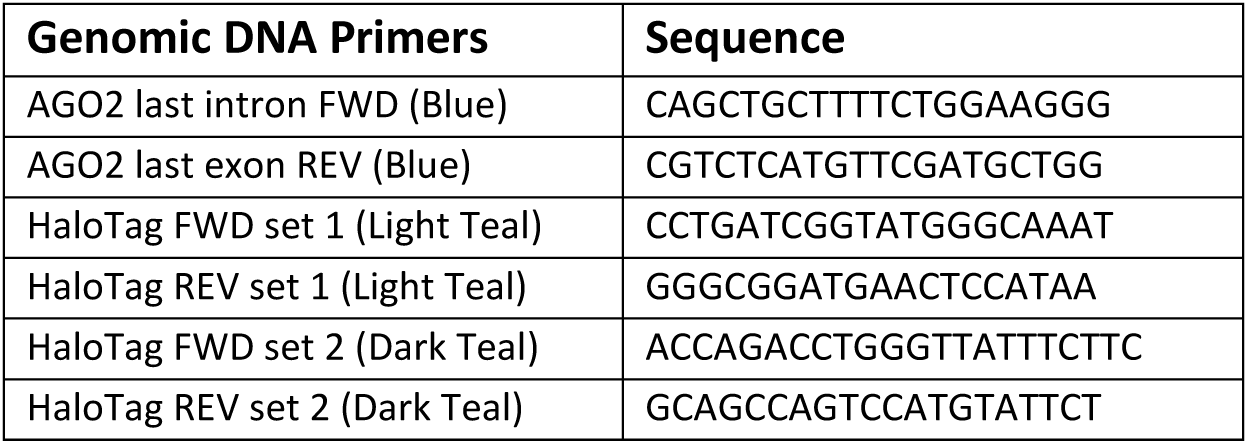

### AGO2 knockout A549 generation

Using the Horizon Discovery Edit-R CRISPR system, wild type A549 cells were transfected with Cas9 nuclease mRNA, TRACrRNA and two crRNAs targeting AGO2 at exon 2 and exon 15 according to manufacturer’s instructions. DharmaFECT Duo transfection reagent was used to transfect cells. Four days post-transfection, some cells were collected to extract genomic DNA for T7 endonuclease assay to assess editing efficiency and the rest of the cells were expanded. From the heterogenous cell population, single cell clones were obtained by plating cells at 0.7 cells per well in 96 well plates. Single cell clones were screened for loss of AGO2 expression by immunoblotting using rabbit anti-AGO2 (Sino Biological 101620-T36). AGO2 knockout clones were expanded and frozen in liquid nitrogen.

### Cloning of AGO2 in pEGFP-C1 and pEGFP-N1

AGO2 cDNA was generated from total human RNA from H358M cells using a gene-specific reverse primer and the SuperScriptIII reverse transcriptase (Life Technologies), according to manufacturer’s instructions. Using 2X Q5 hot start PCR master mix (NEB), AGO2 cDNA was amplified from the cDNA using forward and reverse primers containing specific restriction enzyme sites for the vectors to be cloned into: for cloning into pEGFP-C1 (to make N-terminally tagged AGO2), HindIII in the forward primer and BamHI in the reverse primer; for cloning into pEGFP-N1 (to make C-terminally tagged AGO2), EcoRI in the forward primer and BamHI in the reverse primer. Following agarose gel electrophoresis to confirm correct size of products, gel extraction was carried out and restriction digestion performed on insert and vector DNA. Ligation was performed using Instant sticky-end ligase mix (NEB) and DH5a E. coli were transformed with ligation reactions. Bacteria were selected on LB agar plates containing kanamycin. Colonies were picked and miniprep cultures were set up. Miniprepped DNA (prepared with Monarch plasmid DNA extraction kit) was sequenced to confirm insertion of AGO2 at the correct sites.

To generate a vector expressing WT AGO2 without a tag, pEGFP-C1 AGO2 was digested with NheI-HF and HindIII to excise the EGFP cassette. This digestion was run on an agarose gel and the band for the remaining vector was excised and gel extracted. Sticky ends were blunted with blunt enzyme mix (NEB) and the vector was recirularised using Quick ligase. Following bacterial transformation and selection on LB agar plates containing kanamycin, miniprep cultures were set up and plasmid DNA was extracted and sequenced.

### Analysis of Cell Doubling Time

Cell doubling time was measured using the IncuCyte ZOOM live-cell imaging platform (Essen Bioscience). Cells imaged using the IncuCyte were seeded in triplicate into 24 well plates at 12,000 cells per well and imaged at 10x magnification every 12 hours for 108 hours. Cell confluence per well was calculated using the IncuCyte ZOOM software (Essen Bioscience). Data represent mean ± SEM of three independent experimental repeats.

### Western Blotting Analysis and Densitometry

Cells were lysed in RIPA buffer (150 nM NaCl, 1% (v/v) IGEPAL, 0.5% (w/v) deoxycholic acid, 0.1% (w/v) SDS, 50 mM Tris) supplemented with protease and phosphatase inhibitors (Pierce, Roche) during the log phase of growth. Protein concentration was quantified using the Pierce BCA protein assay kit (ThermoFisher Scientific, 23225). Lysates were electrophoresed on polyacrylamide gels transferred onto PVDF membranes and probed using the following antibodies: mouse anti-beta-Actin (Sigma #A1978), mouse anti-Vinculin (Invitrogen, 14-9777-82), rabbit anti-AGO1 (CST, 5053), rabbit anti-AGO2 (CST, 2897), rabbit anti-AGO2 (Sino Biological 101620-T36), rabbit anti-AGO3 (abcam ab154884), rabbit anti-AGO4 (CST, 6913), mouse anti-HaloTag (Promega, G921A), rabbit anti-DDX6 (Bethyl, A300-460A), rabbit anti-GW182 (Bethyl, A302-329A) (TNRC6A), mouse anti-Dicer (Abcam, ab14601), mouse anti-alpha-Tubulin (GeneTex, GTX628802) and mouse anti-Histone H3 (Upstate, 05-499). Anti-IgG horseradish peroxidase (Dako) and chemiluminescent detection (ThermoFisher Scientific, Millipore WBKLS0500) were used to develop immunoblots. Blots were imaged on a GE Healthcare ImageQuant imager. Densitometry of immunoblot band intensity was performed on ImageJ, with target band intensity normalised to relative loading control intensity. Data represent mean ± SEM of three independent experimental repeats.

### Protein Co-Immunoprecipitation

Cells from a 15 cm dish (one per elution) were lysed in 1 mL NP-40 buffer (150 mM NaCl, 50 mM Tris pH 8.0, 0.7% NP-40, 5% Glycerol) supplemented with protease and phosphatase inhibitors before centrifugation at 14,000 g for 5 minutes at 4 °C to clear. After 80 µL of cleared lysate was taken for input and mixed with 20 µL of 5X Laemmli buffer and boiled for 5 minutes, cleared lysate (1 mL) was mixed with 1 µg of relevant antibody (Mouse IgA isotype control (eBioscience Invitrogen 14-4762-81) and Mouse IgA anti-AGO2 (Santa-Cruz, sc-53521) and rotated at room temperature for 3 hours. Protein L (Pierce, 88849) beads were washed twice in 500 µL of NP-40 buffer, resuspended in 30 µL of NP-40 buffer, and then added to each IP. Samples were conjugated by rotating for 1.5 hours at room temperature. Magnet bound samples were washed 4 times in 1 mL PBS with 0.1% Tween followed by elution of protein complexes by addition of 40 µL of 0.2 M Glycine (pH 2.5) and 3 minutes incubation at room temperature. Elutions were transferred to new tubes and neutralised with 5 µL of 1 M Tris-HCl (pH 8). Following addition of 11.25 µL of 5X Laemmli buffer and boiling for 5 minutes, samples were ready for analysis by western blot as described above.

### Cell Fractionation

Nuclear and cytoplasmic cell fractions were prepared as performed previously.^44^ Following lysis of cells with hypotonic lysis buffer, cells were incubated on ice for 10 minutes and centrifuged at 500 g for 5 minutes at 4 °C. After supernatant was collected to form the cytoplasmic fraction, cell pellets were washed for three minutes in 0.5 mL of hypotonic buffer three times. Nuclear membrane was lysed with nuclear lysis buffer and sonicated for 20 seconds, followed by centrifugation at 15,000 g for 10 minutes at 4 °C, with this supernatant forming the nuclear fraction. Samples were then prepared and analysed by Western blot as described above.

### Luciferase Reporter Assays

nFor Firefly luciferase (FFLuc) silencing assays, cells were plated in 96 well plates. The following day cells were transfected using Attractene reagent (Qiagen, 301005) with 20ng/well pGL3 FFLuc, 10 ng/well pRLSV40 RLuc (expressing Renilla luciferase for normalisation), and 25 ng/well of esiRNA targeting either GFP (control) (Sigma, EHUEGFP) Scr control non-targeting pool siRNA (Horizon Discovery), or esiFFLuc (Sigma, EHUFLUC). After 48h, cells were harvested in 1X Passive Lysis Buffer and l and stored at −80°C. After thawing, FireFly and Renilla Luciferase activities were measured using the Dual-Luciferase Reporter Assay System (Promega, E1960) according to the manufacturer’s instructions on a FLUOstar^®^ Omega microplate reader. Firefly luciferase activity was divided by the Renilla luciferase activity to normalise the data.

For assaying miR-100 silencing activity, we used the psiCheck2 miR-100 Targeting (T) and non-targeting (NT) reporters described in Bridge et al., 2017 (cite the cell reports 2017 paper here). Cells in 96 well plates were transfected with 20 ng of reporter plasmid, 20ng of AGO2 expression plasmid (or empty vector pEGFP-C1), and 15 nM of miR-100 microRNA mimic (Sigma) using Attractene transfection reagent (QIAGEN). 24 hours post-transfection, cells were harvested in 1X Passive Lysis buffer and luciferase activities assayed as above. Renilla luciferase activity was normalised by dividing by the Firefly luciferase activity.

### MiR-451 RT-qPCR

TRIzol reagent (Invitrogen, 15596026) was used to isolate total RNA from cells at indicated confluency (log-phase = ∼60% confluency; confluent = ∼95% confluency) 60 hours after plating according to the manufacturer’s instructions (with an additional 70% ethanol wash of RNA pellets). cDNA was generated from total RNA using the miRCURY LNA RT Kit (QIAGEN, 339340) and then amplified in duplicate by qPCR using the miRCURY LNA SYBR Green PCR Kit (QIAGEN, 339345) on a QuantStudio™ 5 Real-Time PCR machine (Applied Biosystems) with primers targeting *miR-451a* (QIAGEN, 3624799) and *U6* snRNA (v2) (QIAGEN, 3608295). *MiR-451a* CT values were normalized to *U6* and relative (to relevant WT) fold transcript abundance was calculated using the 2^−ΔΔCt^ method. Data represent mean ± SEM of two independent experimental repeats. To assay pri-miR-451, 1 µg of total RNA was treated with amplification grade DNase I (Sigma) according to the manufacturer’s instructions. Diluted RNA was assayed using the Go-Taq 1-Step RT-qPCR system (Promega) for pri-miR-451 (Forward primer: GATGTACTAGTCCGGGCACC Reverse primer: TTGTGACAGGCTGAGCAGAG). *Beta Actin* mRNA was also assayed as a housekeeping gene using primers Forward: CTGGAACGGTGAAGGTGACG and Reverse: AAGGGACTTCCTGTAACAATGCA.

### AGO2 WT RT-qPCR

To assay for untagged *AGO2* mRNA in our cells, total RNA was first extracted using TRIzol reagent according to the manufacturer’s instructions. 1 µg of RNA was treated with amplification grade DNase I (Sigma) according to the manufacturer’s instructions. A six-point serial dilution was prepared of RNA for generation of standard curves. Diluted RNA was assayed for *AGO2* transcript using the GoTaq 1-Step RT-qPCR System (Promega A6020) with primers CTGGCTCCAGGGGACAAG (Forward primer) and CCACTCGGTACACAATCGCT (Reverse primer). *Beta Actin* mRNA was also assayed as a housekeeping gene to normalise AGO2 transcript abundance. Primers were purchased from Integrated DNA Technologies.

### Fluorescence Microscopy

Small round coverslips (1.5 thickness) in a 24 well plate were coated with sterile 0.1% gelatine and then AGO2^-/-^ A549 cells were plated. The following day 400 ng of plasmid (EGFP-VO, EGFP-AGO2 or AGO2-EGFP) was transfected per well using DharmaFECT kb reagent (Horizon Discovery), according to the manufacturer’s instructions. 48h post-transfection, cells were fixed with 4% formaldehyde solution in PBS for 10 minutes. Fixative was washed off with PBS three times and coverslips were mounted onto a drop of mounting media containing DAPI (EverBrite™ Hardset Mounting Medium, (biotium (#23004)) on glass slides, and cured in the dark overnight in preparation for imaging by microscopy. For cells which underwent covalent protein labelling with HaloTag-TMR, prior to fixation HaloTag® TMRDirect^TM^ Ligand (Promega, G2991) was added to cell culture medium to a concentration of 100 nM and incubated overnight. Cells were washed three times in fresh cell culture medium and incubated for 30 minutes before a final wash, fixation and mounting. Slides were imaged with the ZEISS LSM 880 with Airyscan confocal microscope (64 X oil objective).

### Structural Analysis

Atomic resolution structures of AGO2 were accessed via RCSB (https://www.rcsb.org). AlphaFold^45^ predictions of full-length AGO2 were taken from the DeepMind-EBI database (https://alphafold.ebi.ac.uk; Access date: 08/2023). The structure of AGO2^HALO^ was predicted using ColabFold v1.5.3.^46^ Solvent accessible surfaces areas of AGO2 (PDB accession code, 4OLB) were calculated using the POPScomp server (http://popscomp.org:3838/) using a solvent radius of 1.4 Å.^47^ Figures were made using PyMol (Schrodinger).

### Statistical Analysis

Statistical significance was calculated using the Student’s t-test using GraphPad Prism 9.5.0 and is represented as: not significant (ns) > 0.05; *p ≤ 0.05; **p ≤ 0.01; ***p ≤ 0.001 throughout. Data and error bars represent mean ± SEM of three independent experimental repeats, unless otherwise stated. Technical replicates (≥ 2) were performed in each independent experimental repeat for each sample and condition where possible.

## Results

### Creation and Genomic Characterisation of *AGO2^HALO^* Cell Lines

To investigate the functional activity of AGO2 fused at the C-terminus to HaloTag, we first generated *AGO2^HALO^* cells using CRISPaint technology. Following CRISPaint transfection of A549 cells, successful editing should result in the insertion of HaloTag sequence (as well as sequence for a T2A site, Puromycin resistance, and Poly(A) Tail) to the last exon of the endogenous *AGO2* locus (***Fig. 1A***). After transfection, puromycin selection, and initial screening to confirm successful editing to fuse *AGO2* with *HaloTag*, two untagged control (UnTagged *C1* (***UT C1***) and UnTagged *C2* (***UT C2***)) and two *AGO2^HALO^* (***AGO2^HALO^ C5*** *and **AGO2^HALO^ C10***)) clonal cell populations were selected for further evaluation in this study alongside parental ***A549 WT*** cells. By selecting two *AGO2^HALO^* and two UnTagged CRISPaint control lines to compare against each other and to *WT* A549 cells we aspired to accord greater confidence when interpreting results.

Amplification of genomic DNA (gDNA) revealed that, as expected, HaloTag sequence was only present in *AGO2^HALO^* clones (***Fig. 1B***). Sequencing of *AGO2^HALO^ C5* and *C10* cell populations identified two *AGO2^HALO^* variants, one long and one short (***Fig. 1C***). Mutations in the long variant replaced the final two amino acids of AGO2 (FA) before the STOP codon of the *AGO2* sequence, while the shorter variant had lost six amino acids (RTMYFA) before the Linker and *HaloTag* sequences. The two pairs of long and short variant sequences are identical in both *AGO2^HALO^* clones (***Fig. 1D***).^37,38,48^

Curiously, we observed amplification of *WT AGO2* sequence at the *AGO2*-*HaloTag* junction in both *AGO2^HALO^* clones (***Fig. 1B***), indicating retention of non-HaloTagged *AGO2* gDNA in edited *AGO2^HALO^*cells. Sequence alignments of these products identified an additional premature STOP codon (TAA) in both *AGO2^HALO^*clones (***Fig. 1E***). This slightly truncated *AGO2* sequence resulted in appreciable levels of non-HaloTagged *AGO2* mRNA transcript in both *AGO2^HALO^* clones (∼40% of normalised *WT AGO2* mRNA abundance in *A549 WT* cells) (***Fig. 1F***). Crucially, however, we did not observe any of this truncated AGO2 during protein analysis of the lines by immunoblot (even at long exposures and adjusted intensities) (***Fig. 1G***), suggesting the premature STOP codon results in very efficient nonsense-mediated mRNA decay (NMD) of this transcript before translation.^49^ Correspondingly, immunoblot with anti-AGO2 and an anti-HaloTag antibody showed a band in the *AGO2^HALO^* lines at 130 kDa, the correct predicted size for a fusion protein of AGO2 (97 kDa) and HaloTag (33 kDa), (***Fig. 1G-H***), confirming the HaloTag was fused with AGO2 in-frame and translated correctly. For the two UnTagged clones 1 and 2, we performed sequencing of the genomic DNA at the end of the AGO2 coding sequence (***Supp Fig. 1B***) and found that no editing had occurred here, demonstrating that *AGO2* coding sequence can be considered WT in these cells.

### Characterisation of Growth Rate and RISC Abundance in *AGO2^HALO^* Cells

As AGO2 is essential for many cell functions, we first probed whether C-terminal *AGO2^HALO^* resulted in changes to cell proliferation. The observed mean doubling time in *WT* cells of 26.5 ± 2.6 h was broadly similar to that in both UnTagged (*UT C1* = 26.4 ± 0.8 h; *UT C2* = 30.6 ± 2.8 h) and *AGO2^HALO^*cells (*AGO2^HALO^ C5* = 28.4 ± 1.9 h; *AGO2^HALO^ C10* = 27.6 ± 1.8 h), with no statistically significant differences observed and the greatest change (in relation to *WT*) detected in *UT C1* cells (***Fig. 2A***). These data suggest *AGO2^HALO^* fusion does not cause a discernible difference in cell proliferation rates (in this cell line context), and that clonal selection has greater potential than *AGO2^HALO^* fusion to influence cell proliferation rate.

**Figure 2.**
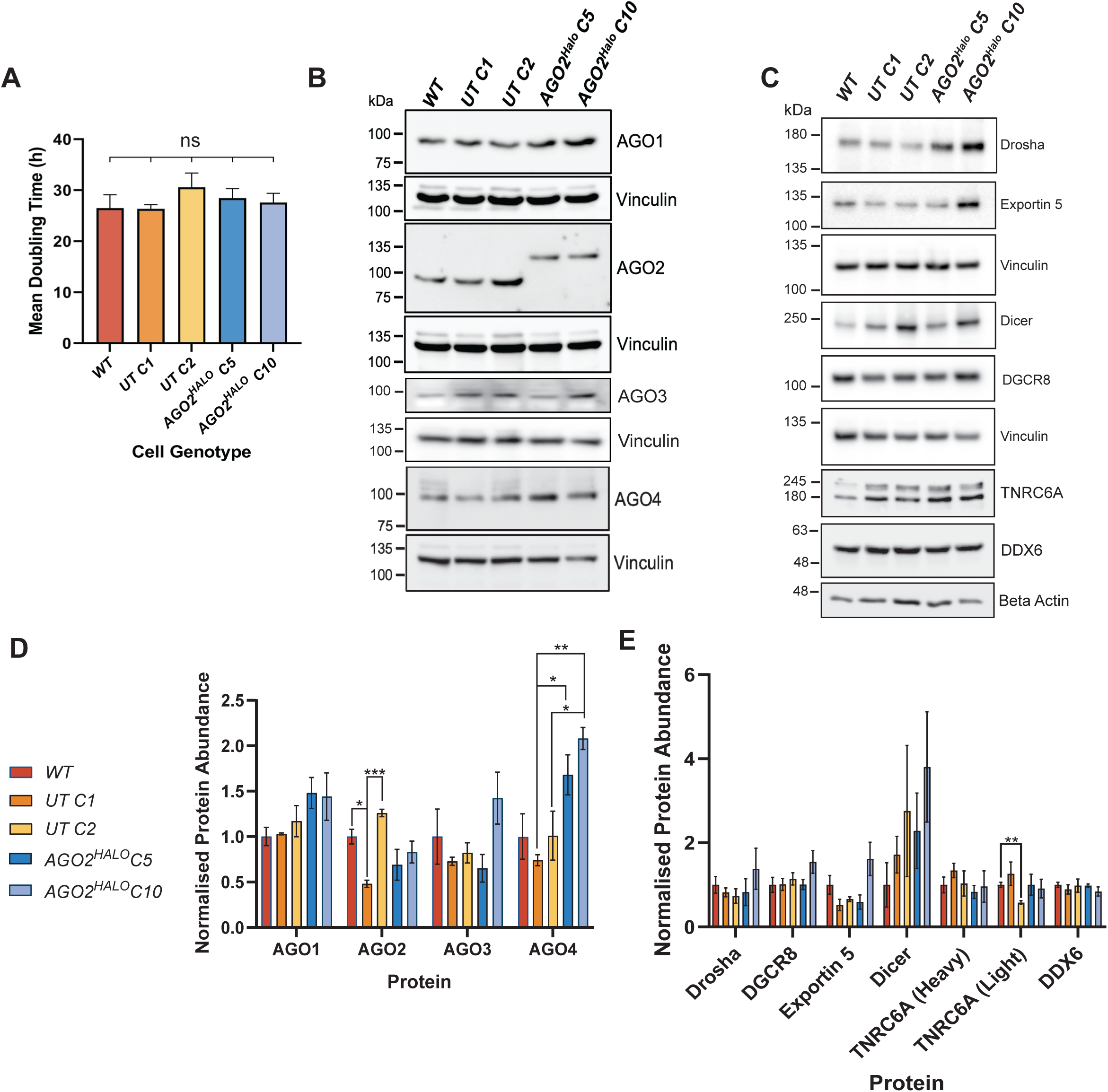
Initial Characterisation of AGO2-HaloTag Cells Lines. (A) Doubling time of A549 WT, two UnTagged (*UT C1* and *UT C2*) and two *AGO2*-*HaloTag* (*AGO2*-*HaloTag C5* and *AGO2*-*HaloTag C10*) measured using IncucyteZoom over 108 hours. Data represent mean ± SEM; n = 3 (ns p > 0.05). (B) Representative Western blot of whole-cell lysates from indicated cell lines (harvested during log-phase) probed with antibodies against AGO1, AGO2, AGO3, AGO4, Vinculin. (C) Representative Western blot of whole-cell lysates from indicated cell lines (harvested during log-phase) probed with antibodies against DDX6, TNRC6A, LIMD1, and Beta Actin. (D) Densitometry of AGO1/2/3/4 (normalised to loading control (Vinculin)) in indicated cell lines. Data represent mean ± SEM; n = 3 (*p ≤ 0.05; **p ≤ 0.01; ***p ≤ 0.001). (E) Densitometry analysis of Drosha, Exportin 5, Dicer, DGCR8, DDX6 andTNRC6A, (normalised to loading control (Beta Actin or vinculin)) in indicated cell lines. Data represent mean ± SEM; n = 3 (*p ≤ 0.05; **p ≤ 0.01; ***p ≤ 0.001).

Due to potential compensatory effects arising from impaired AGO2 function in *AGO2^HALO^* cells, we considered whether the *AGO2^HALO^* fusion resulted in changes in expression of other AGO proteins and other proteins in the miRNA pathway (***Fig. 2B* & *C***). Densitometry of immunoblots (***Fig. 2D* & *E***) showed that both *AGO2^HALO^* clones continued to express other Argonaute family members (AGO1/3/4), the core RISC proteins TNRC6A, DDX6, and proteins necessary for microRNA biogenesis and nuclear export (Drosha, DGCR8, Dicer and Exportin 5). Although the abundance of TNRC6A, DDX6, remained stable across lines, we did note increased AGO1 (not statistically significant), Dicer (not stastistically significant) and AGO4 abundance (statistically significant rise from *WT* and *UT C1* to *AGO2^HALO^ C5* and *C10*) in *AGO2^HALO^* cells compared with WT and UnTagged cells. Speculatively, this could be due to compensation/functional buffering occurring in *AGO2^HALO^* cells to counter impaired AGO2 function.^50^ We also observed a decrease in AGO2 abundance in both *AGO2^HALO^* lines compared with *WT* and *UT C2*, but not *UT C1*, cells. As significant changes in protein abundance were also observed in UnTagged clones compared to *WT*, we were unable to conclude if these decreases were caused directly by *AGO2^HALO^* fusion or clonal selection.

### AGO2^HALO^ Shows Reduced Binding to TNRC6A and Increased Binding to Dicer

A core function of AGO2 is to bind other proteins involved in RNA processing and silencing to form a complex to regulate translational control.^2,6^ We explored whether AGO2^HALO^ can bind TNRC6A - a key protein involved in miRNA-mediated silencing which acts to form a functional RISC when complexed with AGO2^6^ - and Dicer - a critical miRNA processing protein which cleaves precursor miRNAs to generate mature miRNAs and which can also act as a platform for RISC assembly when complexed with AGO2.^51^ We performed immunoprecipitation of AGO2 from WT, UT C1, UT C2, AGO2^HALO^ C5 and AGO2^HALO^C10 cells (***Fig. 3A***). We observed greater levels of Dicer interacting with AGO2^HALO^ than with untagged AGO2. On the other hand, TNRC6A strongly co-immunoprecipitated with AGO2 in WT and UT clones but at reduced levels in the two AGO2-HaloTag clones. Of note, we did observe in elutions for AGO2 immunoprecipitation, a faint band for WT AGO2 at 97 kDa in the AGO2^HALO^ lines, suggesting that some WT AGO2 protein is produced from the AGO2 transcript that avoids nonsense-mediated decay.

**Figure 3.**
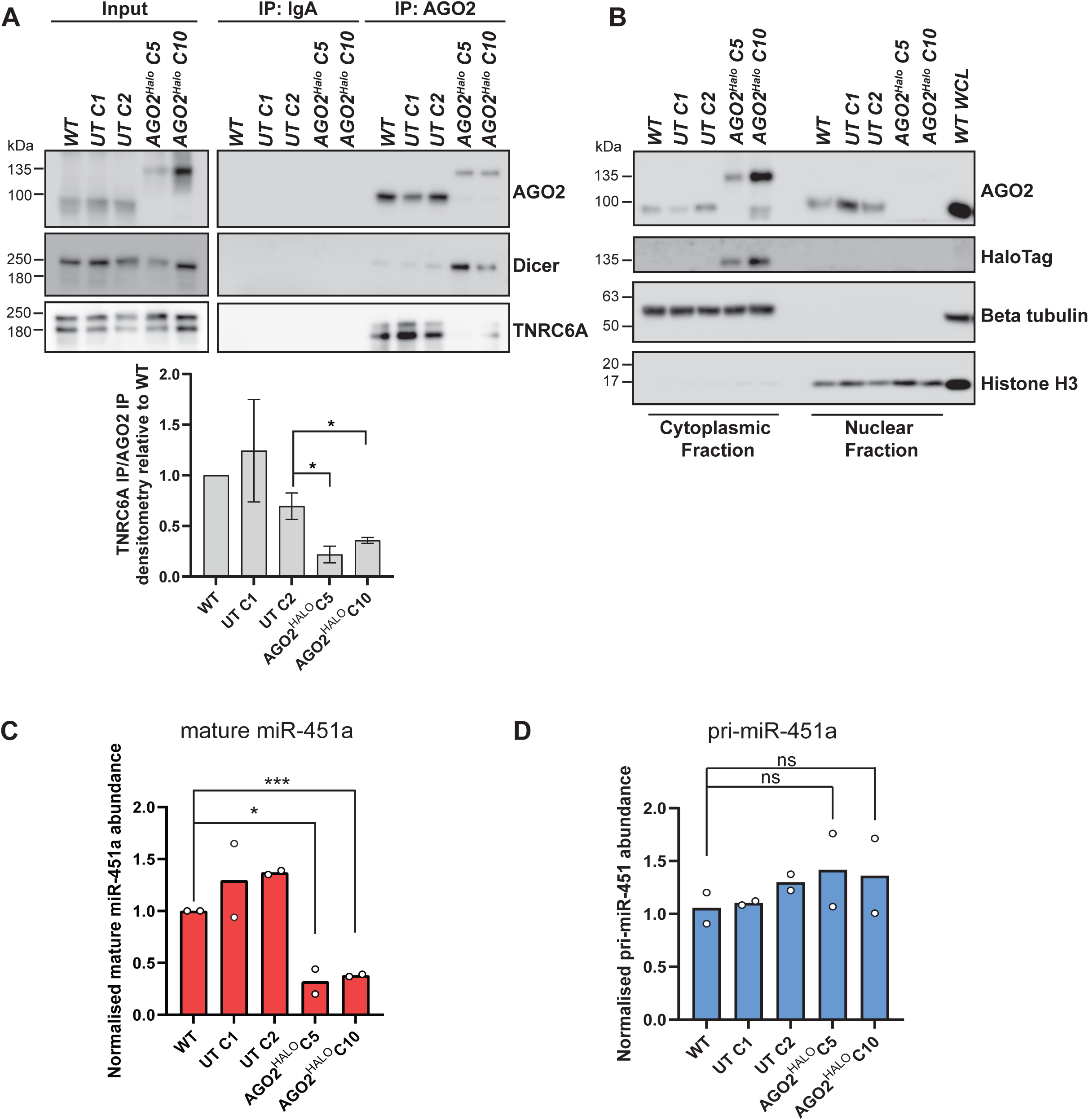
Characterising AGO2 interactions, AGO2 localisation and miR-451a levels in AGO2-HaloTag clones. (A) Endogenous co-immunoprecipitation of AGO2 with Dicer and TNRC6A in WT, UnTagged *C1*, UnTagged *C2, AGO2-HaloTag C5 and AGO2-HaloTag C10* cells. Note the increased AGO2-Dicer and decreased AGO2-TNRC6A co-immunoprecipitation in AGO2^HALO^ cells compared to in WT and UnTagged cells. Histogram shows mean densitometry for TNRC6A IP normalised to AGO2 IP densitometry relative to WT A549 cells. Data represents mean of four independent repeats with * indicating p < 0.05 according to the student’s test when comparing UT C2 to AGO2-HaloTag C5 and C10 lines. (B) Immunoblots of nuclear and cytoplasmic fractions from WT. UnTagged *C1*, UnTagged *C2, AGO2-HaloTag C5 and AGO2-HaloTag C10* cells. Whole cell lysate (WCL) from WT A549 was also blotted alongside. Beta tubulin and histone H3 were probed as loading controls for cytoplasmic and nuclear fractions, respectively. (C) *MiR-451a* abundance (normalized to *U6* RNA and made relative to relevant WT) in indicated cell lines measured by RT-qPCR using the 2^−ΔΔCt^ method. Data represent mean ± SEM; n = 2 (*p ≤ 0.05; ***p ≤ 0.001). (D) Pri-miR-451 abundance (normalised to Beta actin mRNA levels) in indicated cell lines measured by RT-qPCR using the standard curve method. Data represent mean ± SEM; n = 2 (*p ≤ 0.05; ***p ≤ 0.001).

As mRNA translation is regulated temporally and spatially, correct sub-cellular localization of AGO2 and RISC is crucial for normal function. To investigate the sub-cellular localization of the AGO2 fusion protein in *AGO2^HALO^* cells compared with AGO2 in WT and UnTagged cells, we performed nuclear and cytoplasmic fractionation (***Fig. 3B***). Our results show that there is much weaker (almost undetectable at normal exposures) nuclear AGO2 signal in *AGO2^HALO^* cells compared with WT and UnTagged clones. Correspondingly, a clear signal for HaloTag was observed in cytoplasmic fractions of *AGO2^HALO^*cells, but not in the nuclear fraction of these cells.

The pre-eminent function of AGO2 in an active RISC is to correctly and efficiently pair miRNAs with their targets to regulate mRNA translation. This can occur by direct cleavage of the mRNA by AGO2 after small RNA:mRNA binding (with perfect complementarity) or, where (partial or full) seed matching is achieved, through a variety of alternative mechanisms which include recruitment of TNRC6, CCR4-NOT, eIF4F inhibition, ribosome blocking, and mRNA decapping.^2^

To interrogate if *AGO2^HALO^* fusion affected AGO2 cleavage function, we measured miR-451a abundance in our cells. Unlike most miRNAs, maturation of the highly conserved vertebrate miR-451 bypasses Dicer, instead requiring direct cleavage of its precursor hairpin through catalytic AGO2 slicer activity.^52,53,54,55^ Due to this strict AGO2 slicer dependency, the presence of mature miR-451a in the cell can demonstrate AGO2 catalytic function activity. We used RT-qPCR to quantify the normalized abundance of miR-451a in WT, UnTagged, and AGO2^HALO^ cells (***Fig. 3C***). We found mature miR-451a levels to be significantly decreased in the two AGO2*^HALO^* clones when compared to WT and UnTagged clones. By contrast, when we assayed the primary transcript for miR-451 (pri-miR-451), we observed similar levels across all cell lines (***Fig. 3D***), indicating that the differences in mature miR-451a are not due to differences in primary transcript abundance. The reduction in miR-451a abundance compared with UnTagged control cells suggests C-terminal *AGO2^HALO^* fusion resulted in significantly reduced miR-451a biogenesis and therefore impaired AGO2 catalytic activity.

As measures of miR-451a abundance constitute only a qualified test for AGO2 cleavage function, we performed further reporter gene-based functional assays. Using CRISPR-Cas9 editing, we generated AGO2 knockout A549 cells (AGO2^-/-^) to test alongside our WT, UnTagged and AGO2-HaloTag lines (***Fig. 4A***). We more directly assayed AGO2 RNAi/slicer activity with a firefly luciferase (FFLuc) reporter gene that can be targeted by esiRNA (***Fig. 4B***). esiRNA is a heterogenous mixture of siRNAs all of which are fully complementary to the target sequence – FFLuc mRNA in this case. We found that in WT, UT C1 and UT C2 cells transfection of esiFFLuc resulted in a >50% reduction in normalized FFLuc activity compared to where cells were transfected with esiRNA for GFP (a negative control), with no significant difference between these lines. In the AGO2^HALO^C5 and AGO2^HALO^C10 lines, however, no such repression of FFLuc activity was observed, with activity similar to that measured in AGO2^-/-^ cells. This result shows that AGO2 RNAi/slicer activity is completely impaired upon fusion of the HaloTag at the C-terminal of AGO2.

**Figure 4.**
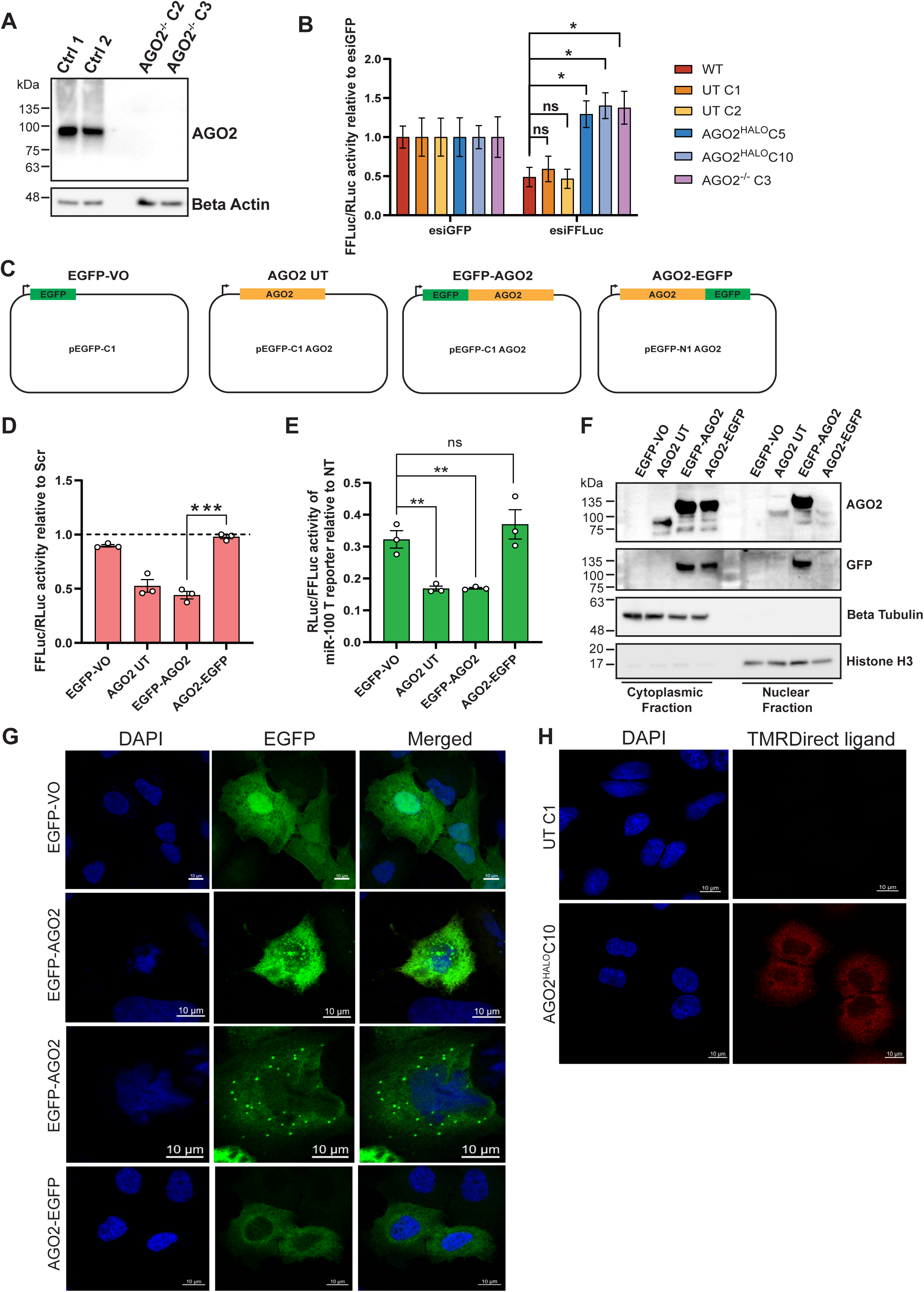
Creation of AGO2 knockout A549, firefly luciferase assays and comparison of N- and C-terminally tagged AGO2. (A) Generation of AGO2 knockout A549 cells. Immunoblot of two AGO2^-/-^ clones alongside two control clones for AGO2 and beta actin as a loading control. (B) Relative Firefly / Renilla Luciferase Activity in indicated cell lines transfected with reporter plasmids expressing Firefly and Renilla luciferase and a siRNA against Firefly Luciferase (esi*FFLuc*) or “non-targeting” control (esi*GFP*). The ratio between Firefly and Renilla luciferase activity was measured 24 h after transfection. Data represent mean ± SEM; n = 3 (ns p > 0.05 according to student’s unpaired t-test). (C) Plasmid constructs for expression of untagged AGO2, EGFP-AGO2 and AGO2-EGFP. EGFP-VO is a control. (D) Relative Firefly / Renilla Luciferase Activity in AGO2^-/-^ A549 transfected with reporter plasmids, AGO2 plasmids and an siRNA against Firefly Luciferase (esi*FFLuc*) or “non-targeting” control (Scr). The ratio between Firefly and Renilla luciferase activity was measured 24 h after transfection. Data represent mean ± SEM; n = 3 (ns p > 0.05; ***p ≤ 0.001 according to student’s unpaired t-test). (E) Relative Renilla/Firefly luciferase activity of a miR-100 targeted (T) reporter relative to a non-targeted (NT) reporter in AGO2^-/-^ A549 transfected with reporter plasmid, 15 nM miR-100 mimic ad the indicated AGO2 constructs or EGFP-VO. Data represent mean ± SEM; n = 3 (ns p > 0.05; ***p ≤ 0.001 according to student’s unpaired t-test). (F) Immunoblots of nuclear and cytoplasmic fractions from AGO2^-/-^ A549 transfected with EGFP-VO, AGO2 UT, EGFP-AGO2 or AGO2-EGFP. Beta tubulin and histone H3 were probed as loading controls for cytoplasmic and nuclear fractions, respectively. (G) Fluorescence microscopy images for AGO2^-/-^ A549 transfected with EGFP-VO, EGFP-AGO2 or AGO2-EGFP. Cell nuclei are stained with DAPI and merged images show DAPI and EGFP signal combined. Scale bars indicate 10 µm. (H) UnTagged C1 and *AGO2-HaloTag C10* cells treated with 100 nM HaloTag-TMR ligand visualized using confocal microscopy.

To avoid any confounding effects of the deletions in AGO2 terminal amino acid residues we found when sequencing the AGO2-HaloTag lines (***Fig. 1C-E***), and to facilitate direct comparison of AGO2 N vs C-terminal tagging, we opted to create plasmid-based AGO2 fusions to EGFP that could be transiently transfected into AGO2^-/-^ cells. We cloned the full length AGO2 CDS into pEGFP-N1 or pEGFP-C1 vectors to create C-terminally or N-terminally EGFP tagged AGO2, respectively. For AGO2-EGFP, the linker between A859 of AGO2 and the M1 of EGFP is ADPPVAT, with no deletions of any AGO2 residues. For EGFP-AGO2, the linker between K238 at the C-terminal of EGFP and M1 of AGO2 is SGLRSRAQAS, with no deletions of any AGO2 residues. We also made a construct expressing untagged AGO2 by excising the EGFP cassette from pEGFP-C1 AGO2. As a control, we used the pEGFP-C1 empty vector which only expresses EGFP (***Fig. 4C***). In AGO2^-/-^ cells, we assessed FFLuc silencing by esiFFLuc where EGFP-VO, AGO2 UT, EGFP-AGO2 or AGO2-EGFP had been co-transfected (***Fig. 4D***). Transfection of AGO2 UT or EGFP-AGO2 yielded approximately 50% repression of FFLuc compared to EGFP-VO transfected cells. AGO2-EGFP transfection, however, failed to cause significant FFLuc repression, indicating an impairment AGO2 RNAi function.

Similar results were obtained when using miR-100 targeted (T) and non-targeted (NT) reporter constructs to assay miR-100 mimic-mediated silencing (***Fig. 4E***): AGO2-EGFP did not cause additional repression of the miR-100-targeted reporter when compared to EGFP-VO transfected cells, whereas AGO2 UT and EGFP-AGO2 did. In EGFP-VO transfected AGO2^-/-^ cells, the level of repression of the T reporter was approximately 70% relative to the NT reporter, most likely due to miR-100 acting via AGO1, 3 or 4 in these cells.

We next examined how untagged, N- and C-terminally tagged AGO2 partitioned in cytoplasmic and nuclear fractions (***Fig. 4F***). All three were detected in cytoplasmic fractions, with the EGFP-tagged proteins also detected by anti-GFP antibody. However, only untagged AGO2 and EGFP-AGO2 were detected in the nuclear fractions, with AGO2-EGFP barely detectable. This partitioning mirrors that observed for the C-terminal AGO2-HaloTag, which was also not detected in nuclear fractions (***Fig. 3B***).

To further investigate tagged AGO2 cellular localisation, we utilized the fluorescence signal of EGFP to performe fluorescence microscopy in AGO2^-/-^ A549 cells that had been transiently transfected with EGFP-VO, EGFP-AGO2 or AGO2-EGFP (***Fig. 4G***). EGFP alone was distributed throughout the cell. For EGFP-AGO2, we observed GFP signal in nuclear areas as well as in cytoplasmic foci that are reminiscent of P-bodies. For AGO2-EGFP however, only a diffuse cytoplasmic signal was present, and no cytoplasmic foci or nuclear localization were seen. Indeed, in imaging experiments on the AGO2-HaloTag lines using the TMRDirect HaloTag ligand we found the signal from this ligand to be exclusively cytoplasmic and diffuse (***Fig. 4H***).

## Discussion

### CRISPaint Generated *AGO2^HALO^* Cells have Comparable miRISC Core Protein Component Abundancies and Cell Viability to WT

We successfully used CRISPaint technology to introduce a C-terminal *HaloTag* to *AGO2* to create *AGO2^HALO^ A549* cells (***Fig. 1***). Two identical variants were identified in both *AGO2^HALO^* clones, suggesting selectivity for these two specific fusion events. We note that the specific sequence outcomes of fusion events are hard to predict due to the potential for bases to be lost during repair. If precise fusion occurs, the CRISPaint-*AGO2* targeting gRNA we employed would fuse AGO2 with a C-terminal HaloTag with the loss of the three terminal amino acid residues of AGO2. WT human AGO2 is 859 residues long. However, we identified two unique variants: “long”, 857 residues; and “short”, 853 residues.

Unexpectedly, we detected a non-HaloTagged AGO2 sequence (covering the gene-tag (*AGO2-HaloTag*) junction) and appreciable levels of WT *AGO2* mRNA transcript in both *AGO2^HALO^* clones. The absence of observable non-HaloTagged AGO2 in *AGO2^HALO^*cells by immunoblot, even at long exposures and high input amounts, initially led us to conclude that these transcripts undergo efficient NMD, presumably because of the two premature STOP codons.^49^ However, we did not rule out that some of this non-HaloTagged *AGO2* transcript may be translated into truncated AGO2 protein, even if only stable for a short period. This was indeed observed later in our study, where we detected a faint ∼97 kDa AGO2 signal in our *AGO2^HALO^* lines by immunoblot where AGO2 had been immunoprecipitated (***Fig. 3A***). NMD has been shown to have a broad range of efficiency in cell populations, with some cells degrading essentially all mRNAs while others escape NMD completely.^56^ It remains plausible that, due to the importance of functional AGO2 in the cell, there is survival selection for *AGO2^HALO^*CRISPaint clones which have less efficient NMD and therefore continue to express (possibly at very low levels) WT (truncated) AGO2 protein. However, this argument is somewhat clouded due to the known role of AGO2/RISC as an inhibitory regulator of NMD itself.^57,58^ Despite the retained expression of very low levels of non-HaloTagged AGO2 in *AGO2^HALO^* lines, we concluded from our immunoblots that the abundance of non-HaloTagged AGO2 protein in *AGO2^HALO^*was so low in comparison to the levels of AGO2 seen in WT and UnTagged cells that it would quickly become saturated. Therefore, this truncated AGO2 would have minimal impact on silencing in the cell and so our experiments evaluating silencing function in *AGO2^HALO^* cells remain valid - especially given that we observed disruption to, rather than retention of, function. The different variants and editing events described herein likely reflect the specific hypotriploid karyotype of A549 cells and^59^ the challenge of creating multiple ‘identical’ edits of AGO2 in this complex and highly mutated genomic context, re-emphasising the need for researchers to comprehensively define their tag-editing at the DNA, RNA, and protein level.

Having validated the successful generation of *AGO2^HALO^* fusion cells, we next sought to define the basic characteristics of the cell. Due to the central nature of AGO2 in normal cell function, tagging of AGO2 may have resulted in differential proliferation. Indeed, down-regulation of *AGO2* has been associated with cell proliferation and apoptosis in prostate cancer.^60^ Encouragingly, our data indicated little change in cell proliferation, with clonal selection potentially having a greater influence on proliferation than *AGO2^HALO^*fusion (***Fig. 2A***).

The first indication that C-terminal fusion of *AGO2^HALO^*caused impairment to normal function was noted from the observed increase in abundance of AGO1 and AGO4 in *AGO2^HALO^* lines compared with WT and UnTagged cells. This may be due to ‘compensation’ between Argonaute proteins following defective AGO2 activity. It has been proposed that there is a degree of functional redundancy between Argonaute proteins, where ablation or silencing of one Argonaute can be ‘compensated’ by other Argonaute proteins.^3^ For example, *AGO2* knock-out can result in increased AGO1 abundance, and vice versa, and AGO3 levels were shown to increase in *AGO1*, *AGO2*, or *AGO1*+*AGO2* knock-out cells: generally, *AGO* knock-out results in increased abundance of the remaining AGO proteins to maintain overall AGO expression at a level near that of *WT*.^50^ If *AGO2^HALO^* fusion were to impair normal AGO2 function, the observed increase in AGO1/4 abundance (***Fig. 2B-C***) may be necessary to compensate for the reduced silencing capacity of AGO2^HALO^. We also considered that some changes may simply be caused by clonal selection, highlighted by the significant reduction in AGO2 abundance in *UT C1* cells.

AGO2-HaloTag, in comparison to WT AGO2, exhibited reduced interaction with TNRC6A upon immunoprecipitation (***Fig. 3A***). This critical binding partner is essential for miRNA mediated gene silencing and normal miRISC function. Structural and interaction studies have shown that AGO2 interacts with tryptophan residues in TNRC6A via three tryptophan-binding pockets in the PIWI domain (***Fig. 5D***).^61,62^ Addition of the HaloTag may have disrupted the folding of the PIWI domain of AGO2 rendering two of three of the Trp-binding pockets unable to interact with the tryptophan residues on TRNC6A, thus impairing the interaction. The three tryptophan binding pockets on the surface of the PIWI domain have been shown to act redundantly in binding tryptophan residues in TNRC6B (PMID: 29576456). However, this study also showed that simultaneous mutation of two pockets reduced interaction with the TRNC6B Ago-binding domain. In our AGO2 IP experiments, we see a *reduced* interaction between AGO2-Halo and TNRC6A, not a total abolition, so it may be that two of the tryptophan-binding pockets are misfolded with the addition of the C-terminal HaloTag.

**Figure 5.**
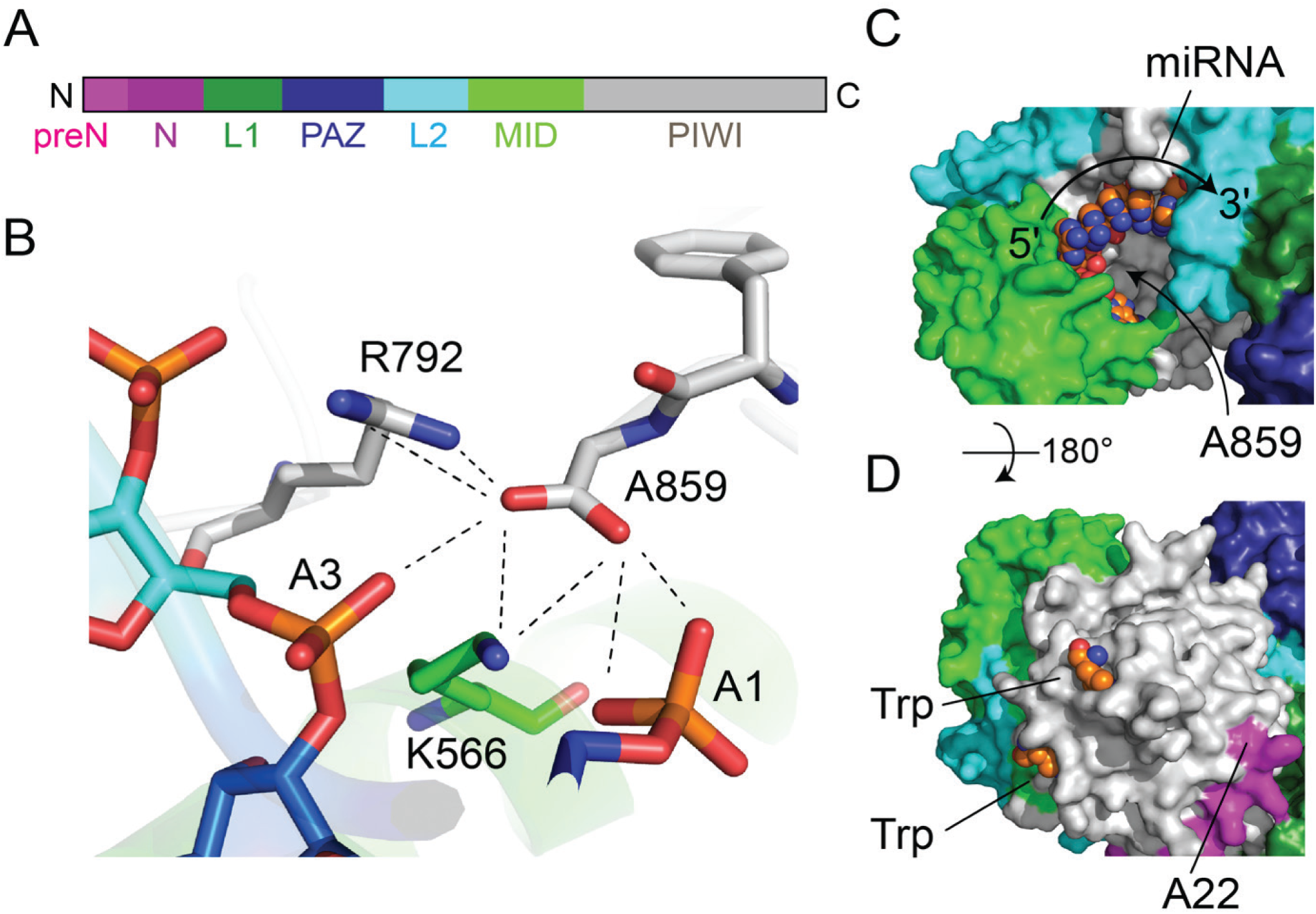
Structural Insights of the C-terminal of AGO2 for an Explanation of Impaired Function Upon Halotag Insertion. (A) Schematic composition of AGO2 showing 7 main domains and motifs. (B) C-terminal residue A859 contributes to miRNA binding (PDB code: 4OLB). Residues shown in stick format and residue type and sequence number annotated. Dashed lines show inter-atom distances < 5.0 Å. (C) Surface representation of AGO2 (4OLB) with domains coloured as in (A). Bound miRNA shown in spheres with 5ʹ-3ʹ direction indicated. The approximate location of the buried C-terminal residue A859 is indicated. (D) Surface representation of AGO2 (4OLB) showing sites of tryptophan binding and the N-terminal most residue (A22) seen in the electron density. Residues 1-21 were not observed in the data.

Co-immunoprecipitation experiments also showed that, compared to WT AGO2, AGO2-HaloTag exhibited greater interaction with Dicer, the protein that is essential for the maturation of almost all miRNAs and the loading of small RNAs on to AGO2. Given the PIWI domain was also shown by others to interact with Dicer,^63^ we can speculate that the structure of the PIWI domain in AGO2-HaloTag still retains the ability to interact with Dicer. Indeed, if TNRC6A and Dicer binding to AGO2 are mutually exclusive, it may be that impairment of TRNC6A interaction leaves AGO2 more available to bind Dicer in *AGO2^HALO^* cells.

### *AGO2^HALO^* Cells Have Significantly Impaired Cleavage and Silencing Function

Ultimately, for C-terminal *AGO2^HALO^* cells to act as a valuable platform to study AGO2 and RISC, data must show that the key protein activities of slicing and miRNA-mediated silencing are sufficiently retained. As such, we focussed on efficiently demonstrating and quantifying catalytic slicing by measuring mature *miR-451a* abundance and RNAi of firefly luciferase in *AGO2^HALO^* cells (***Fig. 3D*; *4B***).

Although the presence of mature *miR-451a* (which strictly relies on AGO2 slicer dependency for maturation) indicates some slicing function in *AGO2^HALO^*cells, the significantly reduced (compared to UnTagged cells) *miR-451a* abundance in *AGO2^HALO^* cells suggests there is consequential deterioration in AGO2 slicing ability arising from *AGO2^HALO^* fusion.

To further explore impairment of AGO2 slicer function, we tested whether an esiRNA for firefly luciferase could silence a firefly luciferase reporter gene transiently transfected into *AGO2^HALO^* cells, alongside *WT*, *UnTagged* and *AGO2^-/-^* cells (***Fig. 4B***). While in *WT* and *UnTagged* cells we observed approximately 50% repression of firefly luciferase expression upon transfection of esiFFLuc, in *AGO2^HALO^*cells no repression of firefly luciferase expression was observed. This was also the case in *AGO2^-/-^* cells, suggesting a near total impairment of AGO2 RNAi function in *AGO2^HALO^* lines.

The *AGO2^HALO^* cell lines we generated using CRISPaint editing and tag insertion led to deletion of some amino acids at the C terminus of AGO2. The results we obtained showing functional impairment of AGO2-HaloTag cannot therefore be specifically attributed to the C-terminal fusion of the tag or the loss of the C -terminal amino acids. Creation and use of the EGFP-tagged plasmid constructs, which resulted in no deletion of residues from AGO2, therefore reduced any ambiguity that C-terminal tagging itself impairs AGO2 function and localisation, as well as providing greater clarity that C-terminal tagging is more deleterious than N-terminal tagging of AGO2. Utilising our *AGO2^-/-^ A549* as a ‘clean’ background, we directly compared the function of N- and C-terminally tagged AGO2 exogenously expressed from transfected plasmid constructs. We used the EGFP tag as it is of similar size to HaloTag and permits straightforward visualization. The firefly luciferase RNAi assay confirmed that a C-terminally tagged AGO2 is impaired in RNAi function (***Fig. 4D***) and we also found that miRNA (miR-100)-mediated silencing is similarly abrogated (***Fig. 4E***). Notably, N-terminally EGFP tagged AGO2 performed as well as an untagged form of AGO2 in these assays, indicating that core AGO2 function is maintained, and demonstrating there is less disruption to AGO2 function when tags are added to the N-rather than C-terminus.

Comparison of our reporter data with previously published evidence of the effect of N-terminal *HaloTag-Ago2* fusion in MEFs – noting the different model systems and editing tools used - indicates that C-terminal *AGO2^HALO^* or EGFP fusion in A549 cells results in greater impairment to silencing function. Dual-Luciferase reporter experiments which were related, but not interchangeable, to ours did not show measurable differences in repression between WT and *Ago2^Halo/Halo^* MEFs for three miRNA binding site containing constructs (*PTEN* 3ʹ UTR *miR-29* binding site; *Adrb2* 3ʹ UTR *let-7* binding site; *Taf7* 3ʹ UTR *miR-21* binding site), though these experiments did not knock-down or perturb active Ago2 and so specific differences between Ago2 and HaloTag-Ago2 activity may not have been assessed.^14^ These findings were re-affirmed by a sensitive two-colour fluorescent reporter system.^14^ Notably, however, the N-terminal *HaloTag-Ago2* fusion did result in notable functional defects including relative derepression of miRNA targets, reduced viability of homozygous mice, and reduced ability to rescue RNAi in *Ago2^-/-^* MEFs compared with WT Ago2.^14^

Notwithstanding, in both comparative and absolute terms our data led us to conclude that C-terminally fused *AGO2^HALO^* cells have significantly impaired endogenous AGO2-mediated silencing function compared to *WT* and *UnTagged* cells. This evidence of such impairment to AGO2 silencing caused by C-terminal fusion of *AGO2^HALO^*does not support C-terminal *AGO2^HALO^* as a suitable model for the study of AGO2 and RISC biology. Regarding the validation of tagged AGO2, we urge for increased use of orthogonal standardised and reproducible (where possible) experimental measures of AGO2 silencing capacity. We believe this will ensure comparability between studies (and tags/tagging strategies), as well as ultimately saving researchers valuable time and resources.

### A Molecular Explanation for Impairment of AGO2 by Addition of a C-terminal HaloTag

Inspection of the 3D structure of human AGO2^51,64^ provides insight as to why a C-terminal HaloTag impairs activity. Protein termini are typically solvent exposed,^65^ however, the C-terminus of AGO2 is an uncommon example where the last residue is buried (only 20% of the surface area of A859 is accessible to a solvent probe with radius 1.4 Å). The buried carboxylate group of A859 also forms functionally relevant interactions (***Fig. 5A***), including hydrogen bonds with several residues (e.g. K556 and R792) that themselves interact with the phosphate groups of residues 1 and 3 of the miRNA. Moreover, in some of the reported structures, the carboxylate groups of Y857 and A859 interact with a buried water molecule that also interacts with the 5ʹ terminal phosphate group of the miRNA. Evidently, modification of the sequence in this region of the protein via deletion of native residues and addition of a HaloTag will likely perturb the RNA binding properties of AGO2 (***Fig. 5B***). Indeed, prediction of the AGO2^HALO^ structure using AlphaFold2 shows the linker region between AGO2 and the HaloTag exiting AGO2 via the miRNA binding site (data not shown). While the confidence (both pLDDT and PAE) for this region of the model is low, this prediction is consistent with our experimental data.

By contrast, the N-terminus of AGO2 is solvent exposed (***Fig. 5D***), which is consistent with previously reported successful tagging of this end of the protein for both *in vitro* and in cell studies. Indeed, crystal structures of AGO2 tend to lack data for the first 20-25 residues,^51,64^ presumably due to the inherent flexibility of this region. Prediction of the structure of full-length AGO2 using AlphaFold2^45^ shows low confidence for the first 25 residues, consistent with this region being flexible and solvent-exposed.^66^

### Distinct Sub-Cellular Localisation of AGO2^HALO^

For proteins to function normally, they must localise correctly. We determined from our imaging and fractionation data that sub-cellular localisation of AGO2 is significantly altered as a result of the C-terminal fusion, with a much-reduced nuclear localisation of the AGO2 fusion protein in *AGO2^HALO^* cells (***Figs 3A and 4H***). Nuclear and cytoplasmic fractionation and imaging experiments also revealed that AGO2-EGFP was similarly absent in the nucleus compared to untagged AGO2 and EGFP-AGO2 (***Fig. 4F* and *G***), suggesting this is an effect specific to C-terminal AGO2 tagging.

Notably, *Ago2*^-/-^ MEFs expressing an N-terminal HaloTag-Ago2 fusion were also found to show a prevalently cytoplasmic localisation of the fusion protein,^14^ though it is unclear if this differs from the signal present in WT cells due to a lack of direct comparison. Impaired nuclear localisation could arise for many reasons, most obviously perhaps due to the addition of the large (33 kDa) HaloTag preventing binding of mediators of nuclear import.^67,68^ It may also be the case that loss of the carboxylate group of A859 specifically disrupts the structure of AGO2 in a way that impairs binding to nuclear import factors. Factors that regulate nuclear import of AGO2 continue to be revealed^69,70^ and it may be that the PIWI domain and/or the C-terminal region are more critical in these interactions than is currently understood. Further studies with deletion mutants of AGO2 would help define the critical residues and structural motifs. Although RISC activity has been considered primarily cytoplasmic, and our principal research intention was to use *AGO2^HALO^* to investigate (primarily) cytoplasmic RISC activity, nuclear RISC has been shown to regulate both transcriptional rates and post-transcriptional mRNA, and RNAi factors are known to be present and functional in human cell nuclei.^71,72,72,73,74^ Moreover, a shift in localization of AGO2 from cytoplasm to nucleus was recently shown to derepress cytoplasmic AGO2-eCLIP targets that were candidates for canonical regulation by miRISC.^75^ Therefore, the potential downstream effects of disrupted nuclear localisation should not be overlooked when considering *AGO2^HALO^*as a model for studying AGO2/RISC.

Previous studies have shown that there is a greater, but comparable, concentration of nuclear and cytoplasmic AGO2 in A549 cells and generally a nuclear to cytoplasmic ratio below one in most cell types.^74,76^ Our fractionation data is congruent with this, with comparable levels of AGO2 observed in both fractions of *A549 WT* cells. Interestingly, tagged AGO2 has previously been shown to result in altered AGO2 protein localisation. N-terminal tagged FLAG-AGO2 exhibited shifted nuclear and cytoplasmic distribution (approximately equal for endogenous AGO2 in T47D cells compared with 67% cytoplasmic and 33% nuclear in *FLAG-AGO2* over-expressing cells),^8^ although this change could be more a consequence of high (5-fold) ectopic overexpression, rather than directly attributable to the tag itself. Similarly, immunofluorescence images of *MYC-AGO2* U2-OS cells stained with FITC-conjugated anti-Myc showed a predominantly cytoplasmic Myc-AGO2 signal with prevalent cytoplasmic foci, while endogenous AGO2 showed a predominantly nuclear signal – it remains unclear if these differences arise directly from the Myc-tag or non-specific antibody binding.^77,78^

Together with our reporter assays indicating significantly impaired silencing function, the distinct sub-cellular localisation of C-terminally tagged AGO2 convinced us to reject both C-terminal *AGO2^HALO^* fusion and C-terminal AGO2-EGFP as a viable tool for study of AGO2/RISC.

## Conclusion

The capacity to investigate the mechanisms behind biological processes and disease has been greatly enhanced by the ability to produce proteins of interest fused to relevant tags. However, every tag added to a protein has the potential to impede functionality, which may invalidate any experimental conclusions. As a result, careful tag design together with reliable and reproducible validation experiments are essential for advancing (miRNA-silencing) research. This is especially important for proteins like AGO2, which often function as part of larger complexes and have numerous and diverse roles and activities. Concerningly, several previous studies that have used tagged forms of AGO2 to investigate AGO2/RISC have not adequately demonstrated, either through published tag validation or the lack thereof, that core functions of tagged AGO2 are sufficiently maintained. As even small modifications can have functional consequences, this shortfall in validation casts doubt on the validity of some of these findings. Additionally, the paucity of AGO2 antibody validation presents a risk of artificial results arising from non-specific binding of antibodies used to probe AGO2. Therefore, to ensure the validity and robustness of future AGO2/RISC investigational findings, we recommend more comprehensive protein-tag (and antibody) validation work to be performed and published in detail alongside investigational findings.

We constructed C-terminal *AGO2^HALO^* fusion cells to investigate if AGO2^HALO^ was a suitable model for studying RISC biology in human cells. Our research revealed that, while AGO2^HALO^ retained some capacity to form native protein-protein interactions, *AGO2^HALO^*cells displayed distinct sub-cellular localization and significantly reduced silencing function compared to normal AGO2. This loss of function led us to conclude that C-terminal fusion of *AGO2^HALO^* was not appropriate for further research into the biology of AGO2/RISC. Moreover, we showed that similar loss of silencing function and aberrant localisation occurs in cells transfected with a C-terminal AGO2-EGFP plasmid, but not N-terminal EGFP-AGO2, suggesting C-terminal tagging of AGO2 in general is inappropriate and N-terminal tagging is more desirable.

While this study establishes important insights into the consequences of C-terminal HaloTag fusion on AGO2 functionality, several limitations should be noted. First, the endogenous tagging data focussed exclusively on the HaloTag — a relatively large (33 kDa) tag — in a single C-terminal configuration, while the N-vs C-terminal AGO2 exogenous tag comparison data is directly relevant for EGFP only (27 kDa). Given AGO2’s complex domain structure and critical roles within RISC, alternative tags or placements may yield different functional outcomes. Exploring smaller tags or N-terminal fusions could potentially mitigate some of the disruptions observed here and is a recommended avenue for future study.

Ultimately, we hope that our work serves as a valuable case study to underscore the importance of careful validation of all core protein competencies of recombinantly tagged proteins. We strongly encourage future research using N-terminal (or any) AGO2 tag fusions to conduct and publish comprehensive validation assays. It is crucial to avoid relying on any single assay and, instead, perform a combination of experiments (e.g., Luciferase reporter assays, *miR-451a*/slicing assays, co-immunoprecipitation/protein-interaction experiments, and localization studies) to validate the range of core protein functionalities, thereby increasing confidence in investigational findings. A community-wide effort to ensure only the most robustly validated reagents and methods are used to characterise AGO2 and RISC biology would enable the miRNA community to gain valuable insights into RNA silencing biology of greater scientific rigour. More broadly, the availability of AlphaFold models for 220 million proteins means that the design of fusion proteins can now be guided by atomic resolution predictions of protein structure in the absence of experimental data. We therefore strongly recommend well-thought-out design and validation strategies to enhance our understanding of the impact of specific tags and tagging strategies on protein function.

## Author contributions

K.M.S., A.F.F.C, A.A, M.J.P, P.G. and T.V.S. designed and performed experiments and analysed the data. A.A and M.A. provide invaluably technical and experimental assistance. All authors contributed to editing and proofreading the manuscript. K.M.S., A.F.F.C, M.J.P., and T.V.S. wrote the manuscript. T.V.S. supervised and managed all research.

## Acknowledgements

We would like to acknowledge the advice and helpful discussions and additional editorial suggestions of this work/investigation from Prof. Dimitris Lagos, Dr Faraz Mardakheh and Dr Paulo S. Ribeiro.

---

**Supplementary Figure 1.**
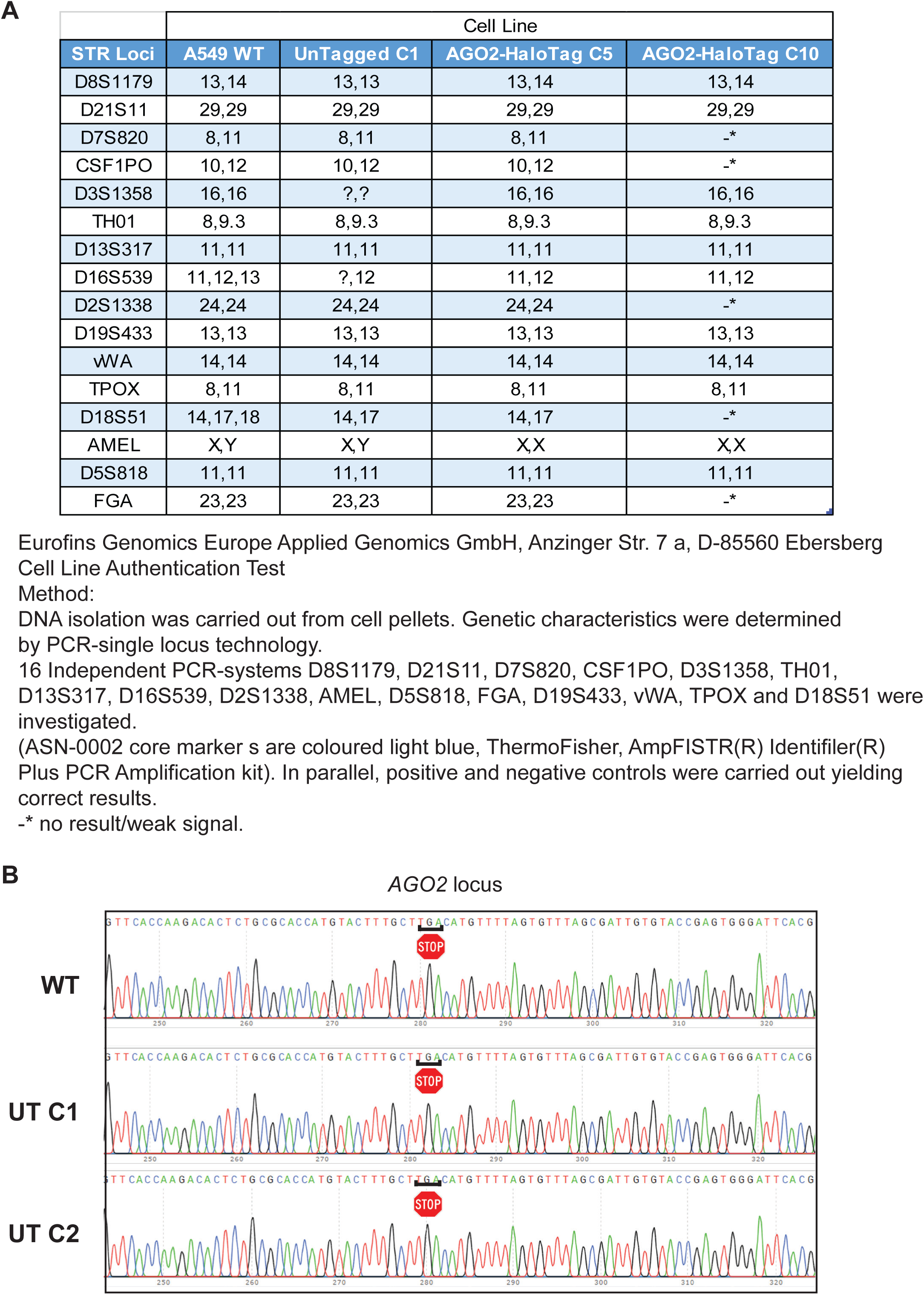
STR Profiling for Cell Line Authentication. (A) Genetic characteristics of indicated cell lines were investigated by STR profiling using 16 independent PCR-systems. Cell Line Authentication Test, Eurofins Genomics Europe Applied Genomics GmbH, Anzinger Str. 7 a, D-85560 Ebersberg Method: DNA isolation carried out from cell pellet (cell layer). Genetic characteristics were determined by PCR-single-locus-technology. 16 independent PCR-systems D8S1179, D21S11, D7S820, CSF1PO, D3S1358, TH01, D13S317, D16S539, D2S1338, AMEL, D5S818, FGA, D19S433, vWA, TPOX and D18S51 were investigated. In parallel, positive and negative controls were carried out, yielding correct results. -* no result / weak signal. (B) Genomic DNA sequencing chromatogram traces of the end of the *AGO2* locus in UnTagged Clone 1 and Clone 2 cells, showing the DNA is not edited. The same region of *AGO2* is shown for WT A549 for comparison.

## References

1. Hutvagner, G. & Simard, M. J. Argonaute proteins: Key players in RNA silencing. Nature Reviews Molecular Cell Biology 9, 22–32 (2008).

2. Frédérick, P. M. & Simard, M. J. Regulation and different functions of the animal microRNA-induced silencing complex. Wiley Interdisciplinary Reviews: RNA 1–22 (2021) doi:10.1002/wrna.1701.

3. Huntzinger, E. & Izaurralde, E. Gene silencing by microRNAs: Contributions of translational repression and mRNA decay. Nature Reviews Genetics 12, 99–110 (2011).

4. Behm-Ansmant, I. et al. mRNA degradation by miRNAs and GW182 requires both CCR4:NOT deadenylase and DCP1:DCP2 decapping complexes. Genes and Development 20, 1885–1898 (2006).

5. Chekulaeva, M. et al. MiRNA repression involves GW182-mediated recruitment of CCR4-NOT through conserved W-containing motifs. Nature Structural and Molecular Biology 18, 1218– 1226 (2011).

6. Lian, S. L. et al. The C-Terminal half of human Ago2 binds to multiple GW-rich regions of GW182 and requires GW182 to mediate silencing. Rna 15, 804–813 (2009).

7. Liu, J. et al. A role for the P-body component GW182 in microRNA function. Nature Cell Biology 7, 1161–1166 (2005).

8. Roya, K. et al. Stable association of RNAi machinery is conserved between the cytoplasm and nucleus of human cells. Rna 22, 1085–1098 (2016).

9. Mauro, M., Berretta, M., Palermo, G., Cavalieri, V. & Rocca, G. La. Special Section on Non-Coding RNAs in Clinical Practice : From Biomarkers to Therapeutic Tools — Minireview The Multiplicity of Argonaute Complexes in Mammalian Cells. THE JOURNAL OF PHARMACOLOGY AND EXPERIMENTAL THERAPEUTICS 52–60 (2023) doi:10.1124.

10. Kedersha, N. & Anderson, P. Mammalian Stress Granules and Processing Bodies. Methods in Enzymology 431, 61–81 (2007).

11. Gibson, T. J., Seiler, M. & Veitia, R. A. The transience of transient overexpression. Nature Methods 10, 715–721 (2013).

12. Booth, W. T. et al. Impact of an N-terminal polyhistidine tag on protein thermal stability. ACS Omega 3, 760–768 (2018).

13. La Rocca, G. et al. In vivo, Argonaute-bound microRNAs exist predominantly in a reservoir of low molecular weight complexes not associated with mRNA. Proceedings of the National Academy of Sciences of the United States of America 112, 767–772 (2015).

14. Li, X. et al. High-Resolution In Vivo Identification of miRNA Targets by Halo-Enhanced Ago2 Pull-Down. Molecular Cell 79, 167–179.e11 (2020).

15. Horman, S. R. et al. Akt-mediated phosphorylation of argonaute 2 downregulates cleavage and upregulates translational repression of MicroRNA targets. Molecular Cell 50, 356–367 (2013).

16. Gurard-levin, Z. A., Kilian, K. A., Kim, J. & Ba, K. HaloTag: A Novel Protein Labeling Technology for Cell Imaging and Protein Analysis. ACS Chemical Biology 60, 45–58 (2010).

17. England, C. G., Luo, H. & Cai, W. HaloTag Technology: A Versatile Platform for Biomedical Applications. Bioconjugate Chemistry 26, 975–986 (2015).

18. Chen, W., Younis, M. H., Zhao, Z. & Cai, W. Recent biomedical advances enabled by HaloTag technology. Biocell 46, 1789–1801 (2022).

19. Gu, J. et al. GoldCLIP: Gel-omitted Ligation-dependent CLIP. Genomics, Proteomics and Bioinformatics 16, 136–143 (2018).

20. Guo, J. K. et al. Denaturing purifications demonstrate that PRC2 and other widely reported chromatin proteins do not appear to bind directly to RNA in vivo. Mol Cell 84, 1271–1289.e12 (2024).

21. Hafner, M. et al. CLIP and complementary methods. Nature Reviews Methods Primers 1, (2021).

22. Grimm, J. B. et al. A general method to improve fluorophores for live-cell and single-molecule microscopy. Nature Methods 12, 244–250 (2015).

23. Duc, H. & Ren, X. Labelling HaloTag Fusion Proteins with HaloTag Ligand in Living Cells. Bio-Protocol 7, 1–8 (2017).

24. Kompa, J. et al. Exchangeable HaloTag Ligands (xHTLs) for multi-modal super-resolution fluorescence microscopy. J. Am. Chem. Soc. 2022.06.20.496706 (2022) doi:10.1021/jacs.2c11969.

25. Kwak, P. B. & Tomari, Y. The N domain of Argonaute drives duplex unwinding during RISC assembly. Nature Structural and Molecular Biology 19, 145–151 (2012).

26. Ma, J. B., Ye, K. & Patel, D. J. Structural basis for overhang-specific small interfering RNA recognition by the PAZ domain. Nature 429, 318–322 (2004).

27. Gu, S., Jin, L., Huang, Y., Zhang, F. & Kay, M. A. Slicing-independent RISC activation requires the argonaute PAZ domain. Current Biology 22, 1536–1542 (2012).

28. Boland, A., Huntzinger, E., Schmidt, S., Izaurralde, E. & Weichenrieder, O. Crystal structure of the MID-PIWI lobe of a eukaryotic argonaute protein. Proceedings of the National Academy of Sciences of the United States of America 108, 10466–10471 (2011).

29. Parker, J. S., Roe, S. M. & Barford, D. Structural insights into mRNA recognition from a PIWI domain-siRNA guide complex. Nature 434, 663–666 (2005).

30. Song, J. J., Smith, S. K., Hannon, G. J. & Joshua-Tor, L. Crystal structure of argonaute and its implications for RISC slicer activity. Science 305, 1434–1437 (2004).

31. Parker, J. S., Roe, S. M. & Barford, D. Crystal structure of a PIWI protein suggests mechanisms for siRNA recognition and slicer activity. EMBO Journal 23, 4727–4737 (2004).

32. Ma, J. B. et al. Structural basis for 5ʹ -end-specific recognition of guide RNA by the A. fulgidus Piwi protein. Nature 434, 666–670 (2005).

33. Rivas, F. V. et al. Purified Argonaute2 and an siRNA form recombinant human RISC. Nature Structural and Molecular Biology 12, 340–349 (2005).

34. Kang, H. W., Tabata, Y. & Ikada, Y. Fabrication of porous gelatin scaffolds for tissue engineering. Biomaterials (1999) doi:10.1016/S0142-9612(99)00036-8.

35. Yuan, Y. R. et al. Crystal structure of A. aeolicus argonaute, a site-specific DNA-guided endoribonuclease, provides insights into RISC-mediated mRNA cleavage. Molecular Cell 19, 405–419 (2005).

36. Nakanishi, K., Weinberg, D. E., Bartel, D. P. & Patel, D. J. Structure of yeast Argonaute with guide RNA. Nature 486, 368–374 (2012).

37. Nakanishi, K. Critical Reviews and Perspectives Anatomy of four human Argonaute proteins. 1–21 (2022).

38. Müller, M., Fazi, F. & Ciaudo, C. Argonaute Proteins: From Structure to Function in Development and Pathological Cell Fate Determination. Frontiers in Cell and Developmental Biology 7, 1–10 (2020).

39. Hauptmann, J. et al. Turning catalytically inactive human Argonaute proteins into active slicer enzymes. Nature Structural and Molecular Biology 20, 814–817 (2013).

40. Schürmann, N., Trabuco, L. G., Bender, C., Russell, R. B. & Grimm, D. Molecular dissection of human Argonaute proteins by DNA shuffling. Nature Structural and Molecular Biology 20, 818–826 (2013).

41. Hur, J. K., Zinchenko, M. K., Djuranovic, S. & Green, R. Regulation of Argonaute slicer activity by guide RNA 3ʹ end interactions with the N-terminal lobe. Journal of Biological Chemistry 288, 7829–7840 (2013).

42. Mukherji, S. et al. MicroRNAs can generate thresholds in target gene expression. Nature Genetics 43, 854–859 (2011).

43. Schmid-Burgk, J. L., Höning, K., Ebert, T. S. & Hornung, V. CRISPaint allows modular base-specific gene tagging using a ligase-4-dependent mechanism. Nature Communications 7, (2016).

44. Liu, Z., Johnson, S. T., Zhang, Z. & Corey, D. R. Expression of TNRC6 (GW182) Proteins Is Not Necessary for Gene Silencing by Fully Complementary RNA Duplexes. Nucleic Acid Therapeutics 29, 323–334 (2019).

45. Jumper, J. et al. Highly accurate protein structure prediction with AlphaFold. Nature 596, 583–589 (2021).

46. Mirdita, M. et al. ColabFold: making protein folding accessible to all. Nature Methods 19, 679–682 (2022).

47. Cavallo, L., Kleinjung, J. & Fraternali, F. POPS: A fast algorithm for solvent accessible surface areas at atomic and residue level. Nucleic Acids Research 31, 3364–3366 (2003).

48. Elkayam, E. et al. The structure of human argonaute-2 in complex with miR-20a. Cell 150, 100–110 (2012).

49. Hug, N., Longman, D. & Cáceres, J. F. Mechanism and regulation of the nonsense-mediated decay pathway. Nucleic Acids Research 44, 1483–1495 (2015).

50. Johnson, K. C., Johnson, S. T., Liu, J. & Chu, Y. Prioritizing Annotated miRNAs : Only a Small Percentage are Candidates for Biological Regulation. (2022).

51. Chendrimada, T. P. et al. TRBP recruits the Dicer complex to Ago2 for microRNA processing and gene silencing. Nature 436, 740–744 (2005).

52. Cheloufi, S., Dos Santos, C. O., Chong, M. M. W. & Hannon, G. J. A dicer-independent miRNA biogenesis pathway that requires Ago catalysis. Nature 465, 584–589 (2010).

53. Cifuentes, D. et al. A novel miRNA processing pathway independent of dicer requires argonaute2 catalytic activity. Science 328, 1694–1698 (2010).

54. Yang, S. et al. Conserved vertebrate mir-451 provides a platform for Dicer-independent, Ago2-mediated microRNA biogenesis. Proceedings of the National Academy of Sciences of the United States of America 107, 15163–15168 (2010).

55. Kretov, D. A. et al. Ago2-Dependent Processing Allows miR-451 to Evade the Global MicroRNA Turnover Elicited during Erythropoiesis. Molecular Cell 78, 317–328.e6 (2020).

56. Sato, H. & Singer, R. H. Cellular variability of nonsense-mediated mRNA decay. Nature Communications 12, 1–12 (2021).

57. Choe, J., Cho, H., Lee, H. C. & Kim, Y. K. MicroRNA/Argonaute 2 regulates nonsense-mediated messenger RNA decay. EMBO Reports 11, 380–386 (2010).

58. Choe, J., Cho, H., Chi, S. G. & Kim, Y. K. Ago2/miRISC-mediated inhibition of CBP80/20-dependent translation and thereby abrogation of nonsense-mediated mRNA decay require the cap-associating activity of Ago2. FEBS Letters 585, 2682–2687 (2011).

59. Park, S. Y., Choi, H. C., Chun, Y. H., Kim, H. & Park, S. H. Characterization of chromosomal aberrations in lung cancer cell lines by cross-species color banding. Cancer Genetics and Cytogenetics 124, 62–70 (2001).

60. Bian, X. J. et al. Down-regulation of Dicer and Ago2 is associated with cell proliferation and apoptosis in prostate cancer. Tumor Biology 35, 11571–11578 (2014).

61. Schirle, N. T. & MacRae, I. J. The crystal structure of human argonaute2. Science (1979) 336, 1037–1040 (2012).

62. Sheu-Gruttadauria, J. & MacRae, I. J. Phase Transitions in the Assembly and Function of Human miRISC. Cell 173, 946–957.e16 (2018).

63. Tahabaz, N. et al. Characterization of the interactions between mammalian PAZ PIWI domain proteins and Dicer. EMBO Rep 5, 189–194 (2004).

64. Schirle, N. T. & MacRae, I. J. The crystal structure of human argonaute2. Science 336, 1037– 1040 (2012).

65. Jacob, E. & Unger, R. A tale of two tails: Why are terminal residues of proteins exposed? Bioinformatics 23, 225–230 (2007).

66. Akdel, M. et al. A structural biology community assessment of AlphaFold2 applications. Nature Structural and Molecular Biology 29, 1056–1067 (2022).

67. Sallis, S., Grondin, B., Leduc, E., Azouz, F. & Pilon, N. FAM172A controls the nuclear import and alternative splicing function of AGO2. bioRxiv (2022).

68. Schraivogel, D. et al. Importin-β facilitates nuclear import of human GW proteins and balances cytoplasmic gene silencing protein levels. Nucleic Acids Research 43, 7447–7461 (2015).

69. Weinmann, L. et al. Importin 8 Is a Gene Silencing Factor that Targets Argonaute Proteins to Distinct mRNAs. Cell 136, 496–507 (2009).

70. Sallis, S. et al. The CHARGE syndrome-associated protein FAM172A controls AGO2 nuclear import. Life science alliance 6, 1–15 (2023).

71. O’Brien, J., Hayder, H., Zayed, Y. & Peng, C. Overview of microRNA biogenesis, mechanisms of actions, and circulation. Frontiers in Endocrinology 9, 1–12 (2018).

72. Nishi, K., Nishi, A., Nagasawa, T. & Ui-Tei, K. Human TNRC6A is an Argonaute-navigator protein for microRNA-mediated gene silencing in the nucleus. Rna 19, 17–35 (2013).

73. Pitchiaya, S., Heinicke, L. A., Park, J. I., Cameron, E. L. & Walter, N. G. Resolving Subcellular miRNA Trafficking and Turnover at Single-Molecule Resolution. Cell Reports 19, 630–642 (2017).

74. Gagnon, K. T., Li, L., Chu, Y., Janowski, B. A. & Corey, D. R. RNAi factors are present and active in human cell nuclei. Cell Reports 6, 211–221 (2014).

75. Johnson, K. C., Kilikevicius, A., Hofman, C., Hu, J. & Liu, Y. Nuclear Localization of Argonaute is affected by Cell Density and May Relieve Repression by microRNAs. (2023).

76. Sarshad, A. A. et al. Argonaute-miRNA Complexes Silence Target mRNAs in the Nucleus of Mammalian Stem Cells. Molecular Cell 71, 1040–1050.e8 (2018).

77. Liu, J., Valencia-Sanchez, M. A., Hannon, G. J. & Parker, R. MicroRNA-dependent localization of targeted mRNAs to mammalian P-bodies. Nature Cell Biology 7, 719–723 (2005).

78. Van Eijl, R. A. P. M., Van Den Brand, T., Nguyen, L. N. & Mulder, K. W. Reactivity of human AGO2 monoclonal antibody 11A9 with the SWI/SNF complex: A case study for rigorously defining antibody selectivity. Scientific Reports 7, 1–11 (2017).

